# Anti-NMDAR and non-anti-NMDAR antibodies promptly modulate NMDAR through p38 and flux-independent signaling: implications for anti-NMDAR encephalitis

**DOI:** 10.1101/2023.06.02.543471

**Authors:** Pavel Montes de Oca-B, Juan Carlos Gómora-García, Lourdes Massieu, Arturo Hernández-Cruz

## Abstract

Anti-N-methyl D-aspartic acid receptor (anti-NMDAR) encephalitis is caused by anti-NMDAR antibodies (Abs) that induce neurologic and psychiatric symptoms, explained mainly by NMDAR hypofunction. In the long-term, these Abs decrease surface NMDAR and NMDAR-mediated intracellular Ca^2+^ ([Ca^2+^]i) influx. However, there are contradictory findings regarding short-term mechanisms. We investigated NMDAR function in cultured neurons after 60 min treatment with three commercial, rabbit, anti-NMDAR Abs (anti-GluN1 extracellular (EC) domain; anti-GluN2B EC domain; and anti-GluN1 intracellular (IC) domain). The anti-GluN2B and anti-GluN1 IC Abs were previously reported to mimic patientś Ab effects in a rodent *in vivo* model and decreased NMDAR-mediated [Ca^2+^]i entry after 24 h treatment in our cells. After 60 min incubation with anti-GluN2B or anti-GluN1 IC decreased the NMDAR-mediated [Ca^2+^]i rise, whereas anti-GluN1 EC slightly increased it. Interestingly, all Abs induced p38 phosphorylation (p-p38). However, surprisingly, it was also elicited by a rabbit Ab directed against a non-NMDAR intracellular epitope, which also reduced NMDAR-mediated [Ca^2+^]i entry. We further investigated the cellular mechanisms regulated by the anti-GluN2B Ab after 60 min. This Ab did not reduce surface NMDAR and p38 inhibition partially prevented its effect on NMDAR function. This Ab did not elicit *per se* an [Ca^2+^]i rise, whereas NMDAR inhibitors 7DCK and MK-801 did not prevent p-p38. Nonetheless, 7DCK prevented NMDAR-mediated [Ca^2+^]i reduction by the Ab, suggesting a role of GluN1 flux-independent signaling. These data indicate that anti-NMDAR and non-anti-NMDAR Ab modulate NMDAR function distinctly and p38 signaling in the short-term, and a role of a third-party mediator. Finally, our results suggest the involvement of NMDAR flux-independent signaling.

## INTRODUCTION

The ionotropic N-methyl D-aspartic acid receptor (NMDAR) plays a critical role in synaptic plasticity mediating long-term potentiation (LTP) and long-term depression (LTD) among other mechanisms (Paoletti, Bellone and Zhou, 2013). Nevertheless, this receptor is widely expressed in different cells and tissues outside the Central Nervous System (CNS), where it plays different roles in other processes that are still under intense investigation (Hogan-Cann and Anderson, 2016). This glutamate ionotropic tetrameric receptor is assembled with two obligate GluN1 subunits, one or two GluN2 subunits (A, B, C or D), and/or one GluN3 subunit (A or B)(Traynelis *et al*., 2010; Paoletti, Bellone and Zhou, 2013; Hansen *et al*., 2018). NMDAR function and intracellular signaling have been studied for decades. It is mainly mediated by Ca^2+^ permeation through its ion channel, which activates different intracellular signaling pathways such as CaMKII, mTOR, MAPK, p38 and calcineurin among others (Hardingham and Bading, 2010; Montes de Oca Balderas and Gonzalez Hernandez, 2018). However, in the last two decades, studies from different groups have unveiled that flux-independent signaling by the NMDAR is also involved in its physiology and physiopathology (Vissel *et al*., 2001; Nong *et al*., 2003; Gérard and Hansson, 2012; Chung, 2013; Tamburri *et al*., 2013; Kessels, Nabavi and Malinow, 2013; Nabavi *et al*., 2013; Stein, Gray and Zito, 2015; Montes de Oca Balderas and Aguilera, 2015; Weilinger *et al*., 2016; Gray, Zito and Hell, 2016; Dore *et al*., 2017; Montes de Oca-B, 2018; Stein *et al*., 2020, 2021; Montes de Oca Balderas *et al*., 2020). Moreover, Kainate, AMPA and GluD glutamate receptors, nicotinic acetylcholine receptors (AChR), voltage-gated calcium channels (VGCC), and voltage-gated potassium channels have also been found to elicit flux-independent signaling (Wang *et al*., 1997; Rodriguez-Moreno and Lerma, 1998; Kaczmarek, 2006; Fernández-Tenorio *et al*., 2011; Cidad *et al*., 2012; Rodrigues and Lerma, 2012; Atlas, 2014, 2022; Valbuena and Lerma, 2016; Jiménez-pérez *et al*., 2016; Richter *et al*., 2016; Zakrzewicz *et al*., 2017; Montes de Oca Balderas *et al*., 2020; Dai *et al*., 2021). Thus, flux-independent signaling by ionotropic receptors seems to be a common mechanism in ionotropic receptors that has been disregarded, as it has been suggested before (Valbuena and Lerma, 2016; Montes de Oca-B, 2018), and as it was concluded in a recent meeting of the American Society for Biochemistry and Molecular Biology (ASBMB) (Montes de Oca-B, 2022).

The NMDAR is involved in different brain pathologies, such as those associated with excitotoxicity, pathological pain, Alzheimeŕs disease, and the recently described anti-NMDAR encephalitis (a-NMDARe) among others (Dalmau *et al*., 2008, 2011; Paoletti, Bellone and Zhou, 2013). The a-NMDARe is an autoimmune disease caused by antibodies (Abs; Ab:antibody) against the NMDAR (anti-NMDAR Abs). The high titers of these Abs in the Cerebrospinal fluid (CSF) of a-NMDARe patients generate neuronal NMDAR hypofunction, resulting in different neurologic and psychiatric symptoms that may lead to hospital admission and even death. These symptoms are reversed when anti-NMDAR Abs titers are reduced with immunotherapy (Dalmau *et al*., 2008, 2011).

Initially, Abs against GluN1/GluN2B heteromers were described in patients of this pathology and it was suggested that the recognized epitopes are conformational, that is, dependent upon their combined conformation (Dalmau *et al*., 2007; Iizuka *et al*., 2008; Tachibana *et al*., 2010; Tabata *et al*., 2014). However, recent findings have hinted that anti-GluN1 Ab are critical for the disease phenotype (Dalmau *et al*., 2008; Gleichman *et al*., 2012; Castillo-Gomez *et al*., 2016). This is intriguing because in a-NMDARe patients with ovarian teratoma the expression of GluN1, GluN2B and other NMDAR subunits has been reported, teratoma associated with >50% of a-NMDARe patients (Tachibana *et al*., 2010; Tabata *et al*., 2014). Given that plasma membrane NMDAR are heteromeric assemblies containing GluN2 subunits, it is possible that the Ab pool in anti-NMDAR patientś includes a mixture of Ab as initially reported and has been suggested, including anti-GluN2B Ab, that could be related to the diversity of neurologic and psychiatric symptoms observed in a-NMDARe patients (Malviya *et al*., 2017). Moreover, anti-GluN2B Abs have been described in other autoimmune diseases (Martinez-Martinez *et al*., 2013).

Some cellular mechanisms that mediate the effects of anti-NMDAR Ab have already been described. Different groups have demonstrated that long-term exposure (>12 h) to anti-NMDARe patientś Ab elicits NMDAR endocytosis, resulting in its hypofunction (Hughes *et al*., 2010; Mikasova *et al*., 2012; Moscato *et al*., 2014; Jézéquel *et al*., 2018). In contrast, apparently contradictory findings have been reported regarding the short-term (<3 h) effects of anti-NMDAR Abs. Some reports have suggested that 1-2 h treatment with patientś CSF or Abs do not decrease surface NMDAR expression in cultured rat hippocampal neurons (Moscato *et al*., 2014; Jézéquel *et al*., 2018). Conversely, other studies report a significant reduction of surface NMDAR after <2 h of incubation with the patient’s CSF (Mikasova et al., 2012; Castillo-Gomez et al., 2016). Moreover, studies at the nanoscale organization of NMDAR found that after two-hour incubation with patientś CSF induced larger clusters of NMDAR with more receptors per cluster (Ladepeche *et al*., 2018). Similarly, different reports account for contradictory findings regarding the short-term effect of patientś Abs on NMDAR function. Mikasova and collaborators also reported diminished intracellular Ca^2+^ signals in response to glutamate following the reduction of cell surface NMDAR (Mikasova *et al*., 2012). Also, after a few minutes of incubation with patientś Abs decreased NMDAR-mediated currents in *Xenopus* oocytes expressing NMDAR (Castillo-Gomez *et al*., 2016). In addition, 5 min treatment with patientś CSF suppressed LTP induction and 1 hour exposure blocked NMDAR-mediated [Ca^2+^]i response in cultured cerebellar neurons (Rubio-Augusti *et al*., 2011; Zhang *et al*., 2012). Also, 1 hour incubation with a patient-derived monoclonal Ab against GluN1 decreased NMDAR-mediated currents (Malviya *et al*., 2017). In contrast, an increase of NMDAR open time has been reported after treatment with patientś CSF (Gleichman *et al*., 2012). Also, no effect was observed on NMDAR-mediated miniature excitatory postsynaptic currents (mEPSC) after 30 min incubation with patientś CSF (Moscato *et al*., 2014), and neither NMDAR-mediated [Ca^2+^]i transients in hippocampal slices were affected after 5 min incubation with patient’s Ab (Jézéquel *et al*., 2018). Thus, there is no consensus regarding the short-term effects and action mechanisms of patientś Abs on NMDAR function, despite they are expected to be critical for the long-term outcome. Moreover, no intracellular signaling that may explain the neuronal response to Abs has been investigated. This lack of consensus could be due to different experimental conditions, such as the diversity of IgG directed against NMDAR subunits present in the Ab pool of patientś samples.

This study analyzed the effects of three commercial, anti-NMDAR, rabbit polyclonal and monoclonal Abs on the NMDAR-mediated [Ca^2+^]i response to short pulses of NMDA/Glycine in cultured rat embryonic cortical neurons. These Abs recognize GluN2B extracellular (EC) domain (anti-GluN2B), GluN1 EC domain (anti-GluN1), or GluN1 intracellular (IC) domain (anti-GluN1 IC)(**Fig. 1**). These Ab were selected because initial approaches to this disease identified Abs against GluN1/GluN2B heteromers in patients (Dalmau *et al*., 2007). For this reason, and the colocalization of patients Ab signal with the commercial anti-GluN2B Ab used here described by this group, in the work of Wurdeman et al. they tested this anti-GluN2B Ab (Wurdemann *et al*., 2016). Importantly, they found that the anti-GluN2B and the anti-GluN1-IC Abs used here mimic the effects of patient’s CSF on LTP depression when stereotactically injected into rat brains. The anti-GluN1EC Ab was selected because it is also a rabbit polyclonal Ab raised against the first 100 aa (out of 540 aa) of the EC N-terminal domain of GluN1 subunit, domain recognized by a-NMDARe patientś Ab (Gleichman *et al*., 2012; Castillo-Gomez *et al*., 2016). We further investigated the cellular mechanisms regulated by the anti-GluN2B after 60 min treatment. Our data indicate that anti-NMDAR and non-anti-NMDAR Abs can modulate NMDAR function in the short-term, in part through p38 signaling, hinting that NMDAR-independent mechanisms, beyond Ab-NMDAR binding are involved. Finally, our results also suggest the involvement of NMDAR flux-independent signaling through the GluN1 subunit.

**Figure 1.**
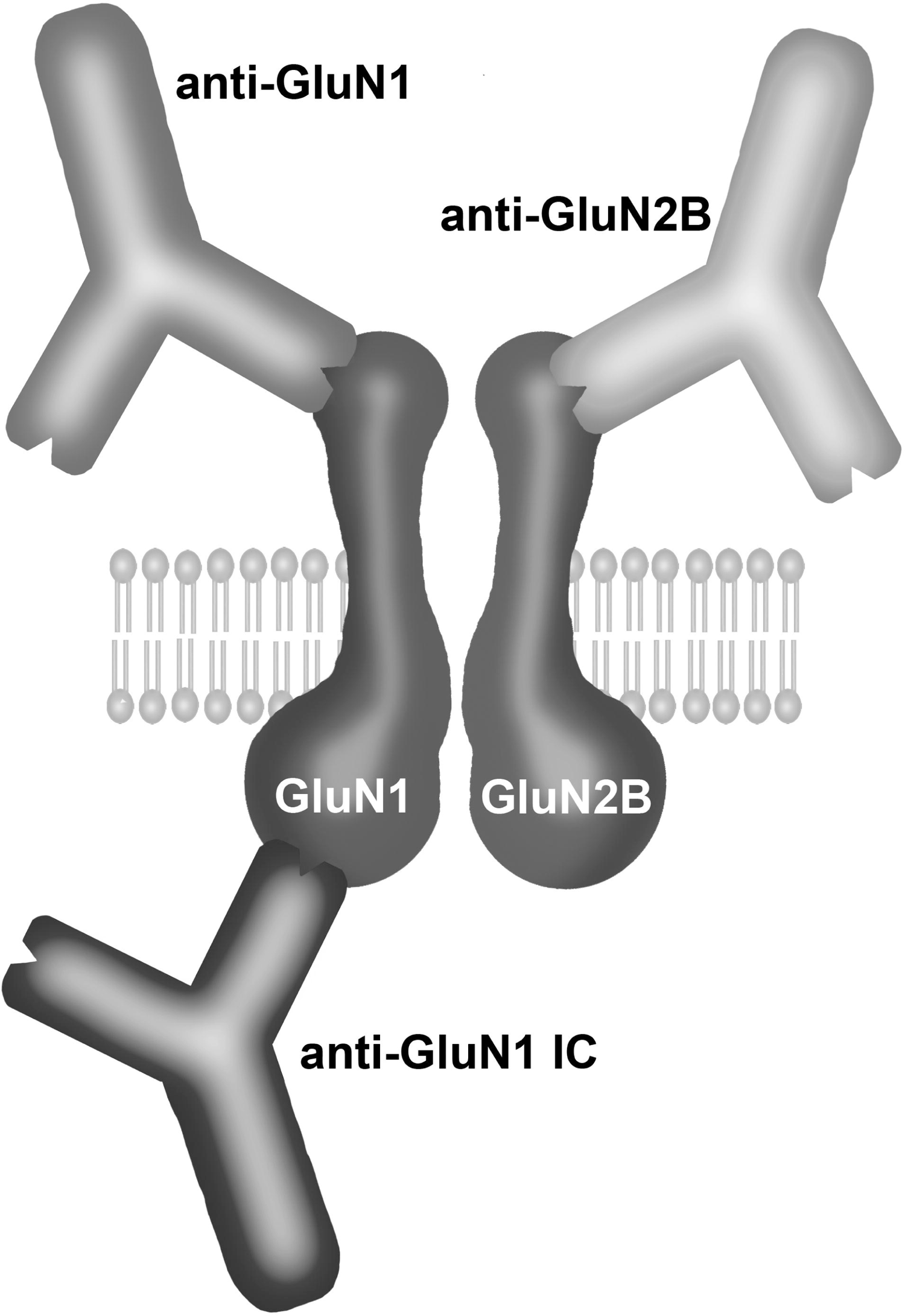
Scheme of anti-NMDAR Ab used in this study. Three different rabbit, anti-NMDAR Ab were used in this study, two against GluN1 subunit, one polyclonal against its extracellular (EC) domain (anti-GluN1, left top Ab) and one monoclonal against its intracellular (IC) domain (anti-GluN1 IC, left bottom Ab). The third one was a polyclonal Ab against the EC domain of GluN2B subunit (anti-GluN2B, top right Ab). Ab were used at 5μg/ml for 24 h or 60 min as described in the text.

## MATERIALS AND METHODS

### Reagents and antibodies

Salts, reagents, NMDA, glycine (Gly), 3-Chloro-4-fluoro-N-[(4-{[2-(phenylcarbonyl)hydrazino]carbonyl}phenyl)methyl]benzenesulfonamide (TCN), α-(4-Hydroxyphenyl)-β-methyl-4-benzyl-1-piperidineethanol (+)-tartrate salt (ifenprodil), and p38 inhibitor SB203580 were from Sigma Chemical Co. (St. Louis, MO, USA). Cell culture media, supplements and Fluo-4 acetomethylester (Fluo-4-AM) were from Life Technologies (Carlsbad, CA, USA). Culture plates were from Costar (Tewksbury, MA, USA). Anti-NMDAR Ab against NMDAR subunits were purchased from Thermo-Scientific (Rockord, IL, USA). These Ab were: rabbit, polyclonal anti-GluN1 against N-terminal EC amino acids 1-100 (cat no. PA5-34599); rabbit, polyclonal anti-GluN2B against N-terminal EC amino acids 1-250 (cat no. 71-8600) and rabbit, monoclonal Ab (MoAb) against the C-terminal intracellular amino acids 834-938 of GluN1 (clone 1H13L3; cat no. 700685). The monoclonal mouse Ab (MoAb) clone R1JHL1 directed against GluN1 EC domain (cat no. PPS011B) used to measure surface NMDAR by immunofluorescence was purchased from R&D Systems (Minneapolis, MN, USA). The “Control Ab” is a rabbit polyclonal Ab against the intracellular domain (C-terminus) of TrKA (sc-118; Santa Cruz Biotechnology, Dallas, TX, USA). The polyclonal chicken Ab directed against MCT2 and the anti-MAP2 Ab were from Millipore (Darnstadt, Germany, AB1287 and MAB378). Secondary Abs were from Jackson Immunoresearch Laboratories (West Grove, PA, USA). NMDAR inhibitor 5,7 dichlorokynurenic acid (7DCK) was purchased from Santa Cruz Biotechnology, (Dallas, TX, USA).

### Animals and Rat Embryonic Cortical Cultured Neurons

Rat Embryonic Cortical Cultured Neurons were obtained as previously described with minor modifications (Gómora-García *et al*., 2021). Briefly, 16-18 days Wistar rat embryos were obtained through cesarean incisions from pregnant rats obtained from the vivarium of the Instituto de Fisiología Celular at the Universidad Nacional Autónoma de Mexico (UNAM) and with approval of the Committee for the Care and Use of Laboratory Animals (CICUAL, LMT160-20). All animal handling and procedures were performed according to the ethical guidelines of the Instituto de Fisiología Celular. Briefly, the cerebral cortex was dissected and chopped, then incubated with 0.25% trypsin/10% EDTA solution at 37 °C for 3 min; the digestion was stopped with a solution containing 0.52% of soybean trypsin inhibitor and 0.08 % of DNase. Cells were suspended in Neurobasal Medium (Gibco, 21103-049, Grand Island, NY, USA) supplemented with 1% of B27 (Gibco, 17504-044), 1% B27 without antioxidants (Gibco, 10889-038), 0.5LmM L-Glutamine, 20 mg/mL gentamycin (Gibco, 15710-064) and plated at a density of 2.2L×L10^5^Lcells/cm2 in plates precoated with poly-L-Lysine. Cells were cultured at 37L°C in a humidified 5% CO2/95% air atmosphere. Cytosine-D-Arabinoside 0.54LmM (Sigma, C-1768) was added to cultures four days after plating. Cells were used for experiments after 8-11 days in *vitro* (DIV).

### [Ca^2+^]i recordings

[Ca^2+^]i determinations were performed as described previously with modifications (Montes de Oca Balderas *et al*., 2020). After incubation with the indicated anti-NMDAR Ab (5μg/ml) and/or drug, cells seeded onto glass coverslips were washed three times and loaded for 35 min with 1 μM Fluo-4-AM in Ringer-HEPES (RH) containing in mM: NaCl 130; KCl 3; CaCl_2_ 2; MgCl_2_ 2; NaH_2_PO_4_ 0.5; NaHCO_3_ 1; HEPES 5; Glucose 5. Cells were then washed three times with RH and placed in the bottom of a plexiglass recording chamber attached to the stage of a custom-made spinning disc confocal microscope (Solamere Technology Group, Salt Lake city, USA) coupled to an upright microscope (Nikon Eclipse 80i; Nikon Corp., Tokyo, Japan) and continuously perfused (1 ml/min) with RH applied to the recording chamber with a syringe injector (New Era Pump Systems Inc; NY, USA). Fluo-4 AM was excited at 488 nm with monochromatic light from a solid-state Coherent Obis laser (Laser Physics, Reliant 100 s488, West Jordan, UT), coupled to a Yokogawa spin-disk confocal scan head (CSUXM1, Yokogawa Electronic Co., Tokyo, Japan and Solamere Technology Group, Salt Lake city, USA). Emitted fluorescence was band-passed (460/50) before collection with a sCMOS (Prime 95B from Teledyne Photometrics, Tucson, AZ, USA). The system was controlled with the open-source microscopy software Micromanager (Edelstein *et al*., 2010). Time-lapse recordings were acquired at 1 Hz. After obtaining baseline recordings for 120 sec in RH, cells were stimulated with a 1-sec pulse of NMDA(50μM)/Gly(10μM) and the recording continued for 120 sec more. NMDA/Gly was applied in RH without Mg^2+^ with a homemade micro-perfusion system consisting of a 0.8 mm syringe tip placed 500 μm above and aside the field of view with an Eppendorf micromanipulator (Eppendorf, Germany), fed by gravity at ≈400 μl/min. Time–lapse recordings were analyzed with Cell Sens Dimension software (Olympus Corporation, Japan). Background noise was subtracted, and circular ROIs of equal size were drawn for each cell soma. The average fluorescence intensity (F) was obtained for each ROI and Ca^2+^ responses were normalized by calculating the relative fluorescence change (ΔF/F_0_) with the formula ΔF/F_0_ = (F*t_x_*-F_0_)/F_0_, where F*t_x_* is F at time *x* and F_0_ is the average basal fluorescence during 100-115 sec of recording, 5 sec previous to the stimulation. ΔF/F was used for comparisons since it abrogates labeling differences.

Selection criteria were applied to exclude cells with a peak response to NMDA/Gly smaller in amplitude to 10% of baseline and those with a bi-spiked response. Data are presented as average response spikes with a range of 80 sec (sec100-180), of cells obtained from at least three different cultures. Each cell response trace was analyzed with a MATHLAB routine that yields the Full Width Half Maxima (FWHM), the spike amplitude and the integrated relative fluorescence change ∫(ΔF/F_0_)(Area Under the Curve, AUC) for each cell, which was calculated as the sum of normalized ΔF/F_0_ values between sec 20 and 50. In addition, raster plots of ΔF/F_0_ vs. time are shown below each spike response for each condition, where the diversity of individual cell responses and the approximate number of cells recorded are appreciated. Some variability was observed in response to NMDA/Gly between cultures, which could be related with their DIV range. For this reason, each recording session included the appropriate control experiments against which the experimental condition was compared. Data of this analysis are given in the text as averages ± SD and in whisker plots, where the *p*-value is also provided.

### Immunofluorescence

Cells seeded onto coverslips were washed with PBS and then fixed with PBS-4% paraformaldehyde (PFA)/4% sucrose on ice (in the case of recorded cells, they were immediately fixed). For some experiments, as indicated, cells were then permeabilized with PBS plus 0.5% Tween-20. After three washes, cells were incubated with the indicated primary Ab (1 μg/ml) for one hour in PBS, washed and then incubated with the appropriate secondary Ab (46.6 ng/ml) for one hour. In some experiments the primary Ab was that of the treatment. Cell nuclei were labeled with Hoechst 33342 (10 μM) for 10 min. Coverslips were then washed and mounted with homemade Mowiol-DABCO. *Z*-stacks were acquired (100X/1.3 N.A. objective) with the spinning disc microscope described above equipped with a motorized stage. Image analysis and quantification were performed using Image J (Schneider, C.A., Rasband and Eliceiri, 2012) as follows: the best-focused slice of the z-stack near the coverslip was extracted and background was subtracted with the Image J built-in sliding ball algorithm. Regions of Interest (ROI) were drawn for each cell soma and images were then segmented with a fixed range. The particles obtained from this segmentation were counted with a cutoff size of 4-infinity pixels to obtain the average density of particles, their area, and average fluorescence intensity. Data of this quantification are presented in whisker plots, where the *p*-value is also given.

### Immunoblotting

Cells were cultured on 12-well cultured dishes. After treatment cells were washed with ice-cold 0.1 M PBS and lysed with 50 μL of buffer lysis (50 mM Tris-HCl pH 8.0,150 mM NaCl, 1% Triton X-100, 0.5% sodium deoxycholate, 1% SDS and 2 mg/mL of protease inhibitor cocktail) per well. Samples were centrifuged at 1500 g at 4°C for 5 min. Protein concentration was determined by Lowry assay and 20 μg of protein from each sample were separated in SDS-PAGE and subsequently transferred to PVDF membranes. The membranes were blocked with 5% dry milk in TBS and incubated overnight at 4 °C with specific monoclonal antibody anti-p-p38 MAPK (Thr180/Tyr182) (1:2000, Cell Signaling, Danvers, MA, USA, 4511) and anti-actin (1:7000, Merck Millipore, Darnstadt, Germany, MAB1501) as loading control after stripping the membrane. A secondary anti-rabbit or anti-mouse antibody coupled to peroxidase was used to detect primary antibodies, and immunoreactivity was detected using the chemiluminescent HRP substrate (Merck Millipore, Darnstadt, Germany). The membranes were exposed using the C-DiGit Scanner (LI-COR, Lincoln, NE, USA).

### Statistical analysis

As described above, statistical differences for the [Ca2+]i responses were evaluated using FWHM, spike amplitude and AUC. These parameters were obtained for each cell response using a MATLAB routine that gets the linear and exponential equations describing the spike of each cell. *p* values for [Ca^2+^]i signals, and immunofluorescence experiments were determined by student’s t-test with Excel (Microsoft). When more than one condition was compared, ANOVA for multiple comparisons test with Tukey correction was performed in GraphPad Prism. In agreement with a recent analysis of the use of statistical *p* value that may mislead data interpretation (Amrhein, Greenland and Mcshane, 2019), no threshold is assigned to claim statistical significance, only the *p* value is given for each comparison.

## RESULTS

### NMDAR-mediated [Ca^2+^]i response in cultured cortical neurons and receptor subunit composition

We first characterized the response of cultured cortical neurons to a brief (1 sec long) pulse of NMDA(50μM)/Gly(10μM) (hereby termed NMDA/Gly) and tested the role and subunit configuration of the NMDAR. For this purpose, we perfused NMDA/Gly in Ringer-Hepes (RH) with or without 2 mM Mg^2+^, since NMDAR channels are blocked by this ion (Hansen *et al*., 2018). **Figure 2A** (left trace) shows NMDA/Gly without Mg^2+^ elicited an [Ca^2+^]i transient. In contrast, the presence of this ion in the NMDA/Gly perfusate strongly reduced the **spike amplitude** (**control**=4.02 ± 1.7 ΔF/F A.U. vs **+Mg^2+^**=0.97 ± 0.82 ΔF/F A.U.), and the **Full Width Half Maxima (FWHM**)(**control**=11.1 ± 4.24 sec vs **+Mg^2+^**=7.06 ± 2.99 sec)(**Fig. 2A** right trace). These kinetic changes resulted in an 83% reduction of the [Ca^2+^]i response (**AUC [Area Under the Curve**]; **control**=50.37 ± 33.05 A.U. vs **+Mg^2+^**=8.17 ± 8.06 A.U.). These data indicate that NMDAR mediates most of the response to NMDA/Gly in our *in vitro* model.

**Figure 2.**
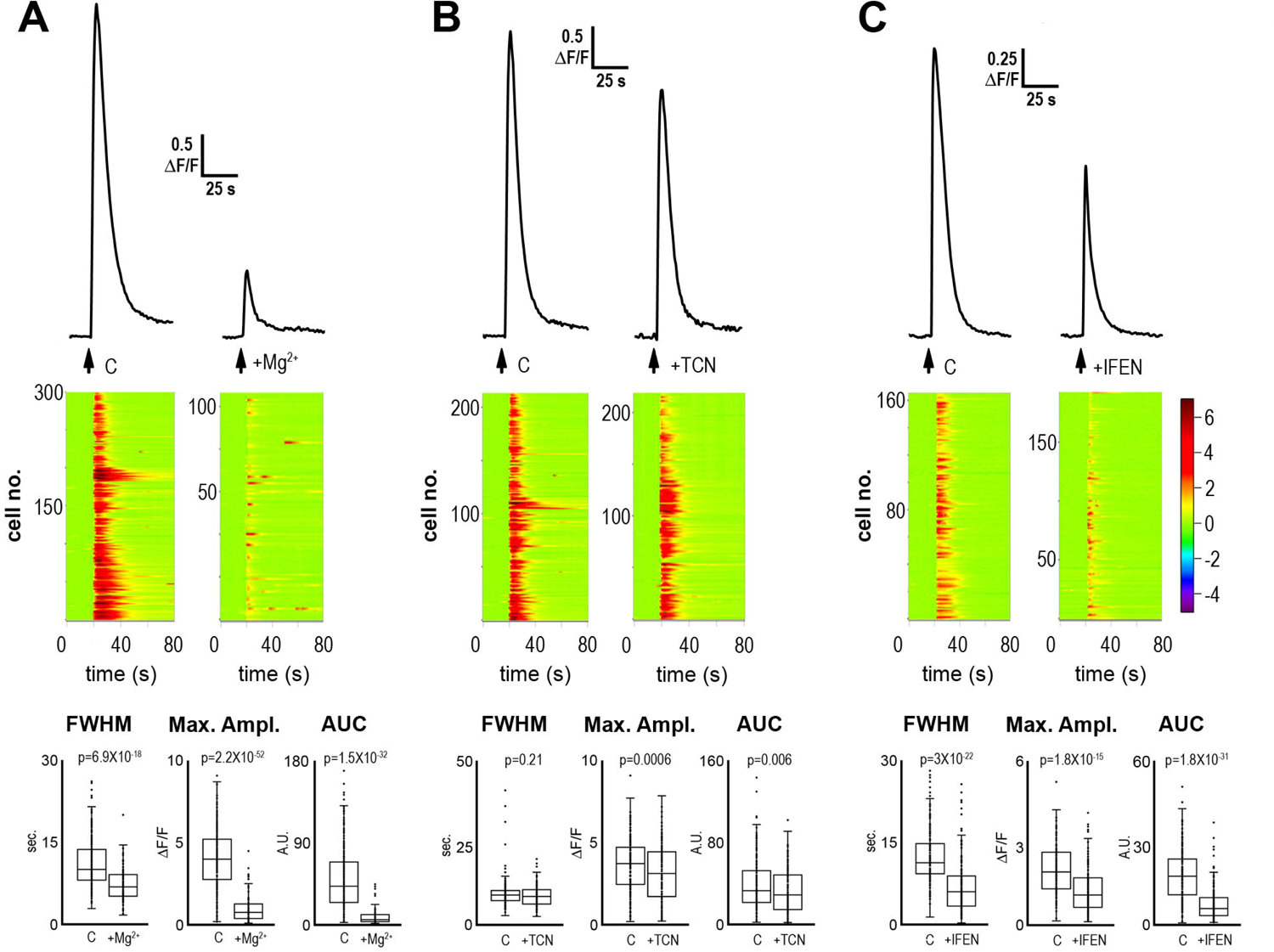
NMDAR-mediated i[Ca^2+^] response and NMDAR configuration. *A* Neuronal averaged response to 1 sec NMDA/Gly stimulation (represented by the arrow below the trace) in 2 mM Mg^2+^ RH (right trace) or Mg^2+^-free RH (left trace). Below each trace, the raster plot of individual cell responses is shown with the same time scale in the *x* axis and the cell number in the *y* axis. The whisker plots for the FWHM, Maximum Amplitude and integrated i[Ca^2+^] (AUC) are shown in the bottom. ***B*** Neuronal averaged response to 1 sec NMDA/Gly stimulation (represented by the arrow below the trace) in control condition (left trace) or with 50 μM of TCN (right trace). Below each trace, the raster plot of individual cell responses is shown with the same time scale in the *x* axis and the cell number in the *y* axis. The whisker plots for the FWHM, Maximum Amplitude and integrated i[Ca^2+^] (AUC) are shown in the bottom. ***C*** Neuronal averaged response to 1 sec NMDA/Gly stimulation (represented by the arrow below the trace) in control condition (left trace) or with 50 μM of Ifenprodil (right trace). Below each trace, the raster plot of individual cell responses is shown with the same time scale in the *x* axis and the cell number in the *y* axis. In the bottom, the whisker plots for the FWHM, Maximum Amplitude and integrated i[Ca^2+^] (AUC) are shown. *p* Student’s t-test values for each comparison are shown above each plot.

We then explored NMDAR subunit composition using two specific inhibitors of NMDAR subunits: TCN, an inhibitor of NMDAR containing GluN2A subunits, and ifenprodil, an inhibitor of NMDAR containing GluN2B subunits. Both inhibitors partially reduced the NMDAR-mediated [Ca^2+^]i response elicited by NMDA/Gly, which is consistent with the presence of both subunits (**Figs. 2B&C**). Nonetheless, ifenprodil inhibited the [Ca^2+^]i response by 57.7 %, whereas TCN inhibited it by 15 %. Also, ifenprodil decreased FWHM and peak amplitude, whereas TCN only decreased the peak amplitude (For TCN ***FWHM*: control**=9.48±4.13 sec. vs **TCN**=8.69±2.91 sec.; ***Max. Ampl*.: control**=3.67±1.56 ΔF/F A.U. vs. **TCN**=3.14±1.64 ΔF/F A.U.; ***AUC*: control**=38.42±24 vs. **TCN**=32.47±21.32. For ifenprodil ***FWHM*: control**=10.31±4.11 sec. vs. **Ifen**=5.82±4.05 sec.; ***Max. Ampl.*: control**=2.01±0.9 ΔF/F A.U. vs. **Ifen**=1.26±0.81 ΔF/F A.U.; ***AUC*: control**=20.92±10.59 vs. **Ifen**=8.85±6.93). These results suggest that at the stage of neuronal maturation of our cultures, GluN2B is more abundant than GluN2A in the NMDAR, although both subunits are expressed.

These experiments demonstrate that [Ca^2+^]i responses to NMDA/Gly in our model of cortical neurons are mediated mainly by NMDAR assembled mostly with GluN2B and less GluN2A subunits, in agreement with what has been described previously in cortical and hippocampal cultured neurons at the same stage of maturation (Li *et al*., 1998; Waxman and Lynch, 2005).

### [Ca^2+^]i responses after exposure to anti-NMDAR Abs

We then tested NMDAR function after 24 h incubation with three different commercial, rabbit, anti-NMDAR Ab against GluN1 or GluN2B subunits (anti-GluN1 extracellular (EC) domain; anti-GluN2B EC domain; and anti-GluN1 intracellular (IC) domain (**Fig. 1**). These Abs were selected for the reasons explained in the introduction at 5 μg/ml, IgG concentration estimated in the CSF of a-NMDARe patients (Jézéquel *et al*., 2018).

Interestingly, after 24h incubation with the anti-GluN1 Ab the total amount of the mobilized [Ca^2+^]i did not change (***AUC*; control**=18.81 ± 13.17 A.U. vs. +Ab a-GluN1=18.28 ± 11.67 A.U.), as it was expected (**Fig. 3A**). This resulted from opposing actions of this Ab on the individual kinetic components of the [Ca^2+^]i response because it reduced the **FWHM** (**control**=8.96 ± 2.42 sec. vs **+Ab a-GluN1**=7.68 ±1.89 sec.) but increased the **peak amplitude** (**control**=1.83 ± 1.17 ΔF/F A.U. vs. **+Ab a-GluN1**=2.1 ± 1.18 ΔF/F A.U.; see **Fig. 2B**). On the other hand, the anti-GluN2B and anti-GluN1 IC Abs decreased the NMDAR-mediated [Ca^2+^]i influx after 24 h treatment in our *in vitro* model, (***AUC*; control**=22.92 ± 14.3 A.U. vs. **+Ab a-GluN2B**=16.32 ± 10.65 A.U.; ***FWHM:* control**=12.46±7.38 sec. vs **+Ab a-GluN2B**= 9.26 ± 5.08 sec.; ***Max. amplitude:* control** =1.88 ± 1.16 ΔF/F A.U. vs. **a-GluN2B Ab** =1.61 ± 0.92 ΔF/F A.U.)(***AUC*; control** = 51.06 ± 33.9 A.U. vs. **+Ab a-GluN1-IC** = 24.46 ± 16.22 A.U.; ***Max. Amplitude*: control**=3.71 ± 1.83 ΔF/F A.U. vs. **+Ab a-GluN1-IC**=2.25 ± 1.32 ΔF/F A.U.; ***FWHM* control** = 12.95 ± 5.09 A.U. vs. **+Ab a-GluN1-IC** = 11.61 ± 5.7 A.U.)(**Figs. 3B &C**).

**Figure 3.**
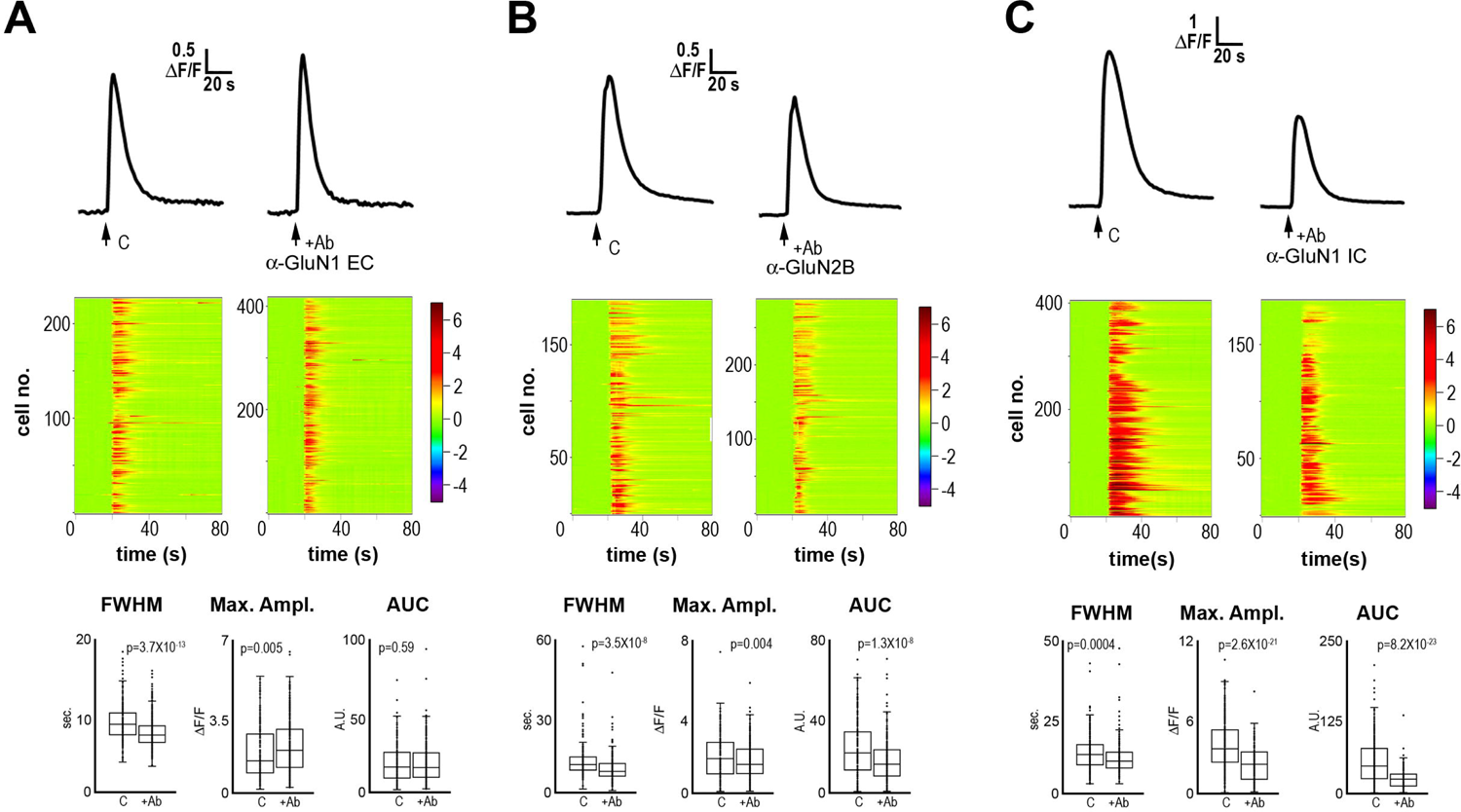
NMDAR-mediated i[Ca^2+^] response after 24h exposure to anti-NMDAR Ab. *A*Neuronal averaged response to 1 sec NMDA/Gly stimulation (represented by the arrow below the trace) in the control condition (left trace) or after 24 h exposure to 5μg/ml anti-GluN1 Ab (right trace). Below each trace, the raster plot of individual cell responses is shown with the same time scale in the *x* axis and the cell number in the *y* axis. The whisker plots for the FWHM, Maximum Amplitude and integrated i[Ca^2+^] (AUC) are shown in the bottom. ***B*** Neuronal averaged response to 1 sec NMDA/Gly stimulation (represented by the arrow below the trace) in control condition (left trace) or after 24 h exposure to 5μg/ml anti-GluN2B Ab (right trace). Below each trace, the raster plot of individual cell responses is shown with the same time scale in the *x* axis and the cell number in the *y* axis. The whisker plots for the FWHM, Maximum Amplitude and integrated i[Ca^2+^] (AUC) are shown in the bottom. The whisker plots for the FWHM, Maximum Amplitude and integrated i[Ca^2+^] (AUC) are shown in the bottom. ***C*** Neuronal averaged response to 1 sec NMDA/Gly stimulation (represented by the arrow below the trace) in the control condition (left trace) or after 24 h exposure to 5μg/ml annti-GluN1 IC Ab (right trace). Below each trace, the raster plot of individual cell responses is shown with the same time scale in the *x* axis and the cell number in the *y* axis. The whisker plots for the FWHM, Maximum Amplitude and integrated i[Ca^2+^] (AUC) are shown in the bottom. *p* Student’s t-test values for each comparison are shown above each plot.

Then, we tested the effect of short-term (60 min) incubation with these Ab. The anti-GluN1 Ab increased slightly the total amount of [Ca^2+^]i mobilized (**AUC; control**=24.4±12.08 A.U. vs. **+Ab a-GluN1**=26.15±11.32 A.U.) (**Fig. 4A**). This effect was achieved through the increase of the **FWHM** (**control**=12.87±4.02 A.U. vs. **+Ab a-GluN1**=15.92±5.02 A.U.) combined with a slight decrease of the **peak amplitude** (**control**=1.84±0.89 ΔF/F A.U. vs. **+Ab a-GluN1**=1.68±0.71 ΔF/F A.U.). On the other hand, the anti-GluN2B Ab reduced the NMDAR-mediated [Ca^2+^]i response (**AUC; control**= 37.78 ± 22.82 A.U. vs. **+Ab a-GluN2B**= 30.05 ± 17.34 A.U.)(**Fig. 4B**), by reducing the **peak amplitude** (**control**=3.68±1.56 ΔF/F A.U. vs. **+Ab a-GluN1**=2.84±1.29 ΔF/F A.U.), without modifying the **FWHM** (**control**=9.08±2.46 A.U. vs. **+Ab a-GluN2B**=9.58±3.08 A.U.). Finally, we found that the anti-GluN1-IC Ab also reduced the mean mobilized [Ca^2+^]i response (**AUC; control**=24.25 ± 17.53 A.U. vs. **+Ab a-GluN1-IC**= 18.08 ± 13.2 A.U.)(**Fig. 4C**). This reduction resulted from decreased **FWHM** (**control** = 10.34 ± 4.92 A.U. vs. **+Ab a-GluN1-IC** = 7.19 ± 3.5 A.U.) In contrast, the peak amplitude did not change (**control** = 2.11 ± 1.3 ΔF/F A.U. vs. **+Ab a-GluN1-IC** = 2.24 ± 1.34 ΔF/F A.U.).

**Figure 4.**
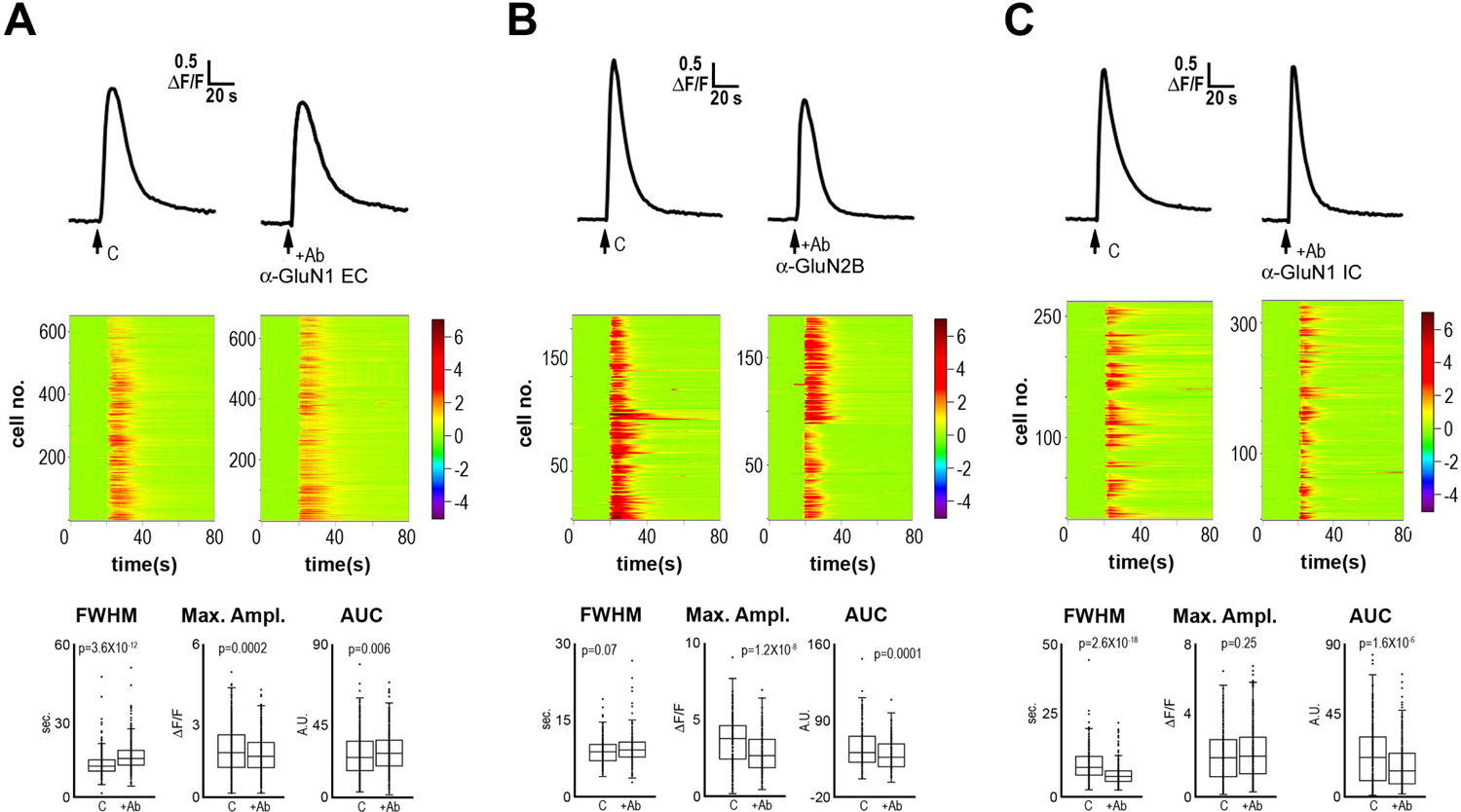
NMDAR-mediated i[Ca^2+^] response after 60 min exposure to anti-NMDAR Ab. *A* Neuronal averaged response to 1 sec NMDA/Gly stimulation (represented by the arrow below the trace) in the control condition (left trace) or after 60 min exposure to 5μg/ml anti-GluN1 Ab (right trace). Below each trace, the raster plot of individual cell responses is shown with the same time scale in the *x* axis and the cell number in the *y* axis. The whisker plots for the FWHM, Maximum Amplitude and integrated i[Ca^2+^] (AUC) are shown in the bottom. ***B*** Neuronal averaged response to 1 sec NMDA/Gly stimulation (represented by the arrow below the trace) in control condition (left trace) or after 60 min exposure to 5μg/ml anti-GluN2B Ab (right trace). Below each trace, the raster plot of individual cell responses is shown with the same time scale in the *x* axis and the cell number in the *y* axis. The whisker plots for the FWHM, Maximum Amplitude and integrated i[Ca^2+^] (AUC) are shown in the bottom. ***C*** Neuronal averaged response to 1 sec NMDA/Gly stimulation (represented by the arrow below the trace) in the control condition (left trace) or after 60 min exposure to 5μg/ml anti-GluN1 IC Ab (right trace). Below each trace, the raster plot of individual cell responses is shown with the same time scale in the *x* axis and the cell number in the *y* axis. The whisker plots for the FWHM, Maximum Amplitude and integrated i[Ca^2+^] (AUC) are shown in the bottom. *p* Student’s t-test values for each comparison are shown above each plot.

These results indicate that depending upon the subunit recognized by the anti-NMDAR Ab, the effects on NMDAR function are distinct. These findings also demonstrate that different mechanisms are involved in the final outcome of NMDAR function after long or short term treatment, since time and peak components of [Ca^2+^]i responses were distinctly regulated by these Abs. Finally, given the early regulation of NMDAR function by these Abs, it is possible that intracellular signaling elicited by them is involved.

### Steric interactions do not mediate the effect of anti-NMDAR Abs

As explained above, the mechanism of NMDAR function regulation by anti-NMDAR Abs in the long-term occurs through endocytosis of surface NMDAR, that in turn leads to NMDAR hypofunction. However, in the short term it has been suggested that the binding of anti-NMDAR Abs to NMDAR may regulate its function through steric mechanisms (Gleichman *et al*., 2012). Therefore, a key question to interpret our results is whether anti-NMDAR Abs remain bound to the receptor during recording, after treatment and Fluo-4 AM labeling. To answer this question, we tested by immunofluorescence the presence of anti-NMDAR Ab in cultured neurons after 60 min Ab treatment, Fluo-4 AM labeling and recording. In these experiments, we immediately fixed the cells after recording. Afterwards, the cells were also incubated with a mouse MoAb against GluN1 to detect surface NMDAR, to then probe this and anti-NMDA Ab used for treatment with appropriate secondary labeled Abs. As shown in **Fig. 5A**, anti-GluN1 and anti-GluN2B Abs used for treatment were not detected on cell surface. Nonetheless, the MoAb revealed surface NMDAR. These findings indicate that anti-NMDAR Abs used for treatment are washed out from the cells during Fluo-4 labeling, washes and recording. Therefore, in contrast with the previous report using a different protocol, in our experiments, steric interaction of NMDAR-Ab seems not to participate in the regulation of NMDAR function when neurons are challenged with NMDA/Gly.

**Figure 5.**
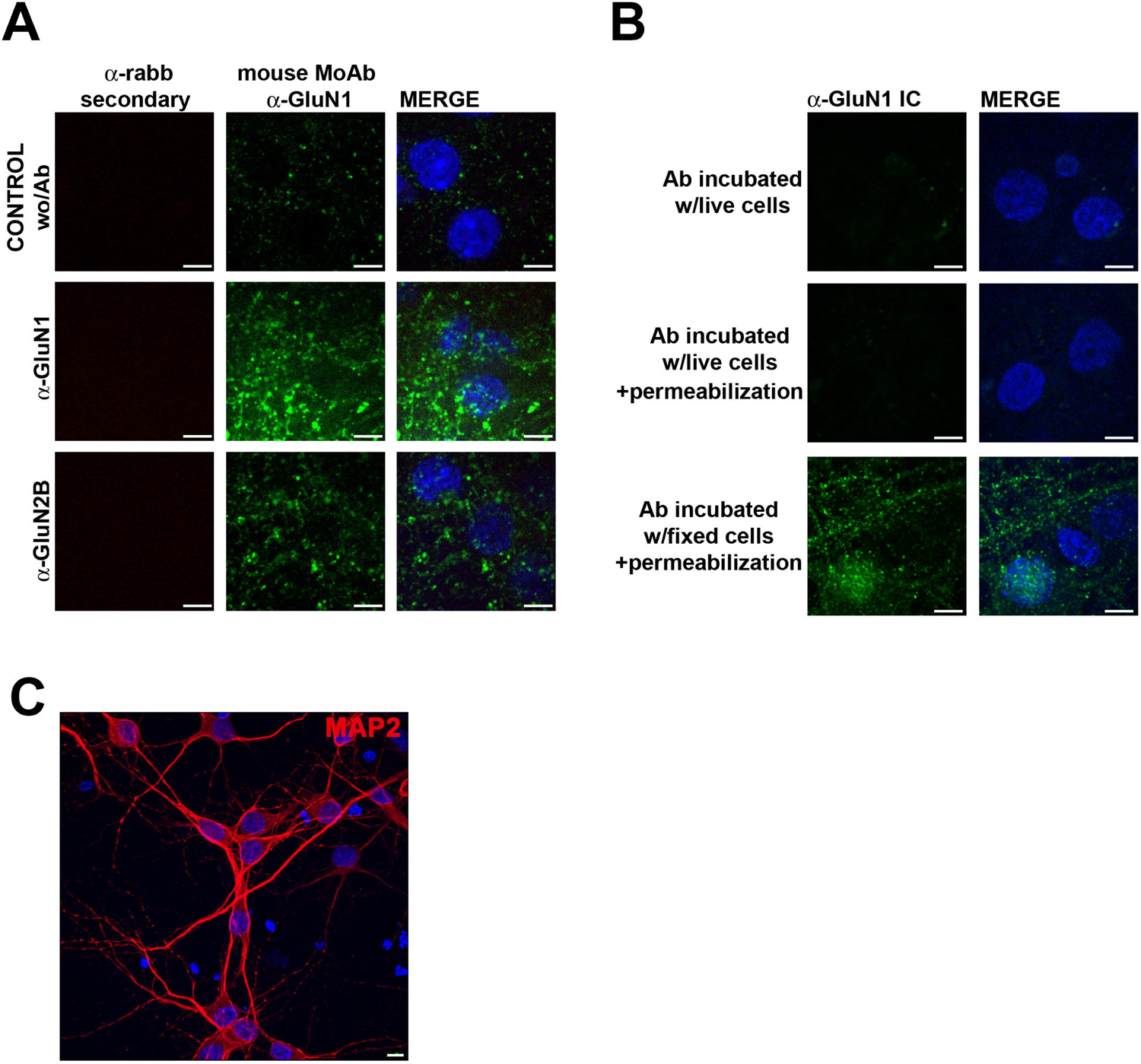
anti-NMDAR Abs do not remain bound to neurons after 60 min treatment, labeling, and recording and anti-GluN1 IC Ab does not access the cell. Representative immunofluorescence images from experiments performed in cultured neurons. *A* Neurons treated or not (control) for 60 min with anti-GluN1 or anti-GluN2B Ab, labeled with Fluo-4 AM, and recorded were fixed to detect the presence of remaining anti-NMDAR Ab with a labeled secondary anti-rabbit Ab. After fixation neurons were first incubated with the mouse MoAb against the GluN1 (clone R1JHL1) to detect surface NMDAR followed by an appropriate secondary labeled anti-mouse Ab *B* Live or fixed neurons were incubated 60 min with anti-GluN1 IC Ab. Then, live cells were fixed and an appropriate secondary Ab was used to detect anti-GluN1 IC Ab on the surface or within the cells (after permeabilization). *C* Representative image of cultured neurons labeled with anti-MAP2 Ab. In the three panels nuclei are shown labeled with Hoechst. Reference bar A & B= 20 μm; C= 10 μm.

We also investigated whether the anti-GuN1 IC Ab can be detected on the neuronal surface after treatment and whether this Ab can access the cell. For these experiments, neurons were incubated with the anti-GluN1 IC Ab for 60 min and then fixed. Some of these cells were permeabilized to detect anti-GluN1 IC Ab that may have accessed the cells. We did not observe staining with a secondary labeled Ab with or without permeabilization. In contrast, abundant labeling was observed when the anti-GluN1 IC Ab was incubated after cell fixation and permeabilization (**Fig. 5B**). These findings show that the anti-GluN1 IC Ab does not remain bound to the cell surface after treatment. Also, they show that this Ab does not access the cell as it has been reported with other Abs (Rhodes and Isenberg, 2017; Slastnikova *et al*., 2018). **Fig 5C** is a representative image of our cultures labeled with MAP2. It can be seen that cells have a typical neuronal phenotype and show MAP2 labeling, with nuclei that are not condensed of fragmented, indicative of healthy neurons, along with some condensed nuclei from cells that degenerated.

### Anti-NMDAR Abs elicit p38 phosphorylation

Given that our experiments suggest that steric interactions between anti-NMDAR Abs and the NMDAR do not have a role, it is then possible that signal transduction elicited by these Abs may be involved. Therefore, we tested if anti-NMDAR Abs could elicit intracellular signaling. For this purpose, we assessed the phosphorylation of MAPK p38 after 60 min incubation with ani-NMDAR Ab, using a standard monoclonal Ab raised against phosphorylated p38 (p-p38). NMDAR agonist and co-agonist binding activate this kinase and more recently, it was found to be involved in flux-independent signaling by the NMDAR (Nabavi *et al*., 2013; Stein, Gray and Zito, 2015; Dore *et al*., 2017; Montes de Oca-B, 2018; Stein *et al*., 2020, 2021). Strikingly, Western Blot experiments showed that the three anti-NMDAR Ab induced p-p38 with different intensities. In contrast, the control condition did not show p-p38 (**Fig. 6**). Also, the anti-GluN2B and anti-GluN1EC Abs induced p-p38 conspicuously compared with control untreated cells. However, the anti-GluN1 IC Ab yielded a robust p-p38 band with a slight upper shift of the molecular size. Intriguingly, the anti-p-p38 Ab also revealed a band of around 53 kDa, which may correspond to a sumoylated form of p38 (Wang *et al*., 2021)(**Supporting Figure 1**). These observations indicate that anti-NMDAR Abs promptly elicit intracellular signaling involving p38 activation. Given the different magnitudes of this phosphorylation, it is suggested that additional mechanisms are involved. In addition, the induction of p-p38 by the anti-GluN1 IC Ab is unexpected because the epitope for this Ab is intracellular, and the Ab would have no access to its epitope. Thus, this finding supports the notion that perhaps other membrane receptors beyond the NMDAR can mediate intracellular signaling in response to these Abs that ultimately regulate NMDAR function.

**Figure 6.**
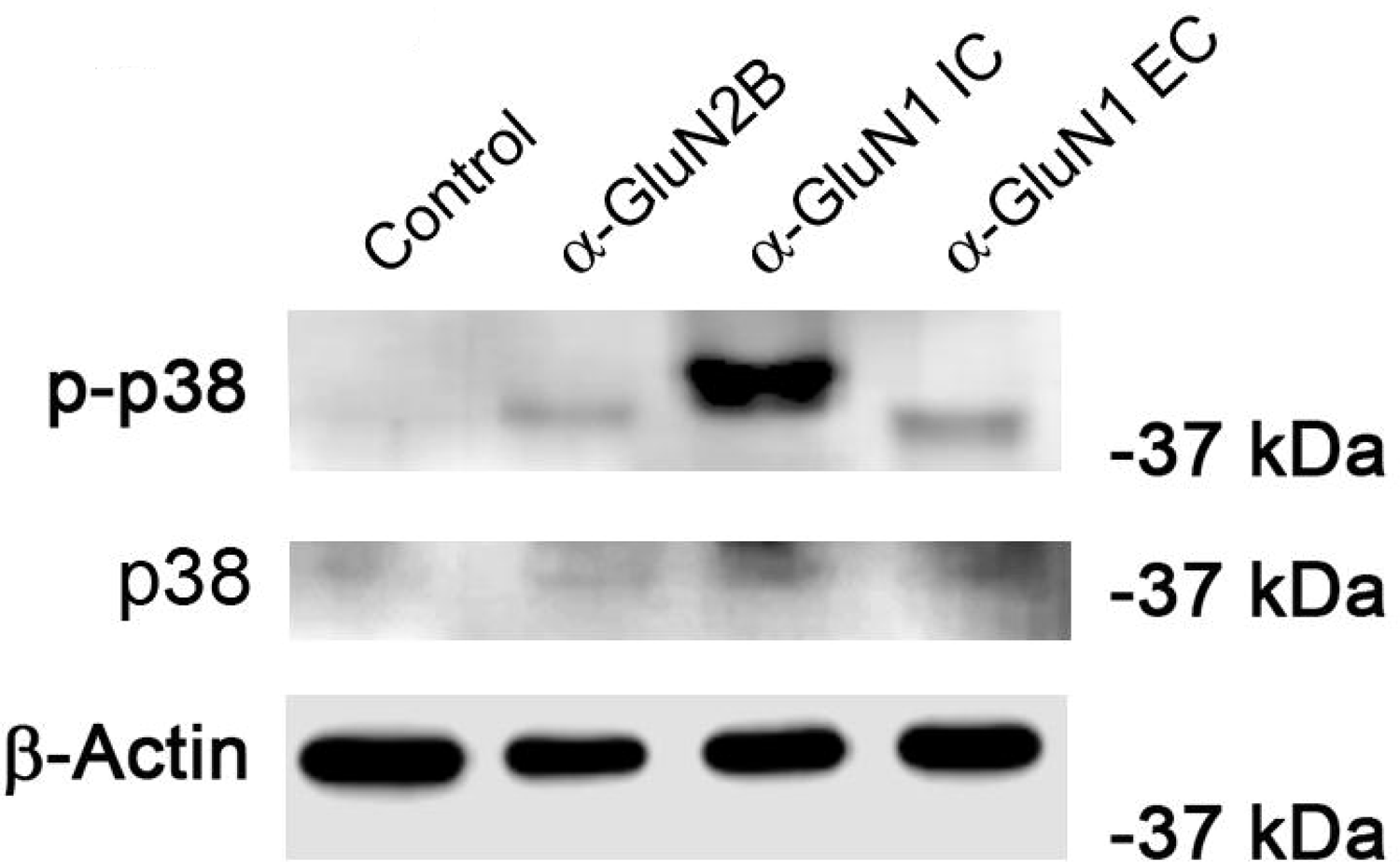
p38 phosphorylation by anti-NMDAR Ab. Western Blot experiment of neuronal lysates treated for 60 min with the three anti-NMDAR Ab or control cells. Total p38 and the loading control (actin) are shown in the panels below for comparison. One representative experiment is shown of at least three independent performed.

### A non-anti-NMDAR Ab also elicits p-p38 and down-regulates NMDAR function

It has been assumed in the literature that the effect of anti-NMDAR Abs on NMDAR function is mediated by Ab binding to the receptor. Therefore, we tested whether other mechanisms unrelated to Ab binding to the NMDAR may be involved in p-p38 induction. For this purpose, we tried a rabbit polyclonal Ab against the intracellular domain of TrkA, an antigen that like the IC domain of GluN1, is not readily accessible to extracellular Abs unless cells are permeabilized. Strikingly, as shown in **Fig. 7A**, after 60 min incubation with this control Ab (C Ab) induced a strong p-p38 band, like that elicited by the anti-GluN1 IC Ab (**Fig. 6**). Importantly, a chicken polyclonal Ab raised against a solute transporter of the cell membrane did not elicit p-p38 (not shown). This finding demonstrates that other rabbit Ab also induce intracellular signaling through the p38 pathway. Therefore, we then tested the effect of this rabbit control Ab on the NMDAR-mediated [Ca^2+^]i response. Unexpectedly, this unrelated control Ab also reduced importantly the total amount of [Ca^2+^]i that permeated the NMDAR, through the regulation of both time and peak components (***FWHM***: **control**=10.67±4.66 sec. vs **C Ab**=9.08±3.6 sec.; ***Max. Ampl*.: control**=2.48±1.37 ΔF/F A.U. vs. **C Ab**=2.12±0.97 ΔF/F A.U.; ***AUC***: **control**=29.62±23.27 vs. **C Ab**=21.16±11.84) (**Fig. 7B**).

**Figure 7.**
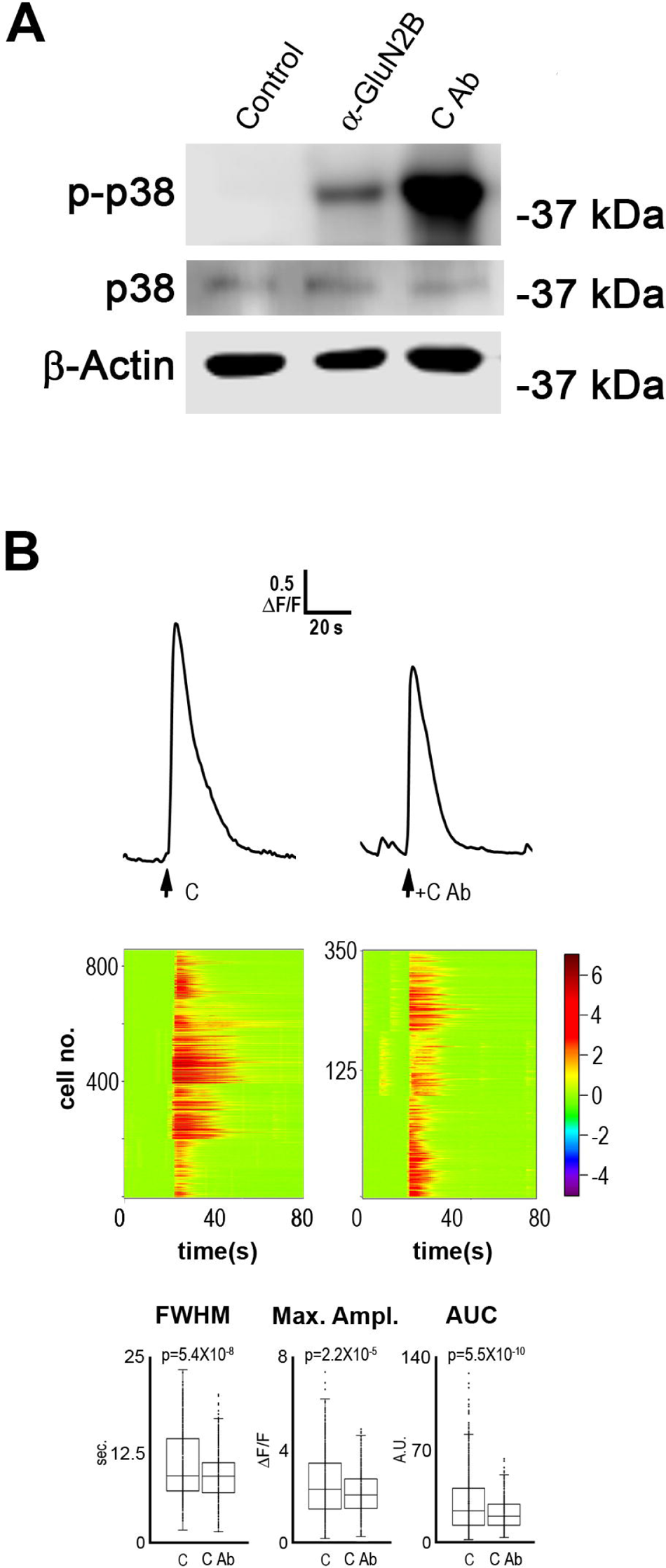
A non-anti-NMDAR Ab induces p38 phosphorylation and down-regulates NMDAR-mediated i[Ca^2+^] response. *A* Western Blot experiment of neuronal lysates treated for 60 min with a control rabbit polyclonal Ab (anti-TrkA IC domain) or control cells. Total p38 and the loading control (actin) are shown in the panels below for comparison. One representative experiment is shown of at least three independent performed. ***B*** Neuronal averaged response to 1 sec NMDA/Gly stimulation (represented by the arrow below the trace) in the control condition (left trace) or after 60 min exposure to 5μg/ml control Ab (C Ab; right trace). Below each trace, the raster plot of individual cell responses is shown with the same time scale in the *x* axis and the cell number in the *y* axis. In the bottom, the whisker plots for the FWHM, Maximum Amplitude and integrated i[Ca^2+^] (AUC) are shown. *p* Student’s t-test values for each comparison are shown above each plot.

Together, these observations demonstrate that p-p38 in our *in vitro* cell model is induced by rabbit, anti-NMDAR directed against extracellular domains, but also by rabbit, anti-NMDAR and non-anti-NMDAR Ab that presumably cannot reach their intended intracellular targets, consistently with idea that only few Abs with specific features are capable of crossing the plasma membrane (Rhodes and Isenberg, 2017; Slastnikova *et al*., 2018). Therefore, these findings strengthen the argument that distinct Abs can regulate NMDAR function through mechanisms other than binding to the NMDAR and reinforce the idea that other cell membrane neuronal molecules could be involved in switching on the p38 pathway.

### Anti-GluN2B Ab does not reduce surface NMDAR and p38 inhibition partially prevents the down-regulation of NMDAR function

We wanted to explore further the mechanisms that modulate NMDAR function after exposure of neurons to anti-NMDAR Abs. We focused our studies on the anti-GluN2B Ab since it recognizes the extracellular domain of the NMDAR, as most Ab detected in a-NMDARe patients, and because it down-regulates NMDAR function not observed with the anti-GluN1 Ab.

As explained above, some groups have reported NMDAR endocytosis after <2 h treatment with patientś Abs, which in turn reduced the NMDAR-mediated [Ca^2+^]i response. Therefore, we measured cell surface NMDAR immunostaining using the MoAb against GluN1, in non-permeabilized neurons after 60 min incubation with the anti-GluN2B Ab. Our experiments showed abundant cell-surface NMDAR but no reduction, as evaluated by the percentage of the cell area covered by NMDAR-specific staining (**control**=6.75±2.73%, n= 32 vs **+Ab a-GluN2B**=8.28±4.08%, n=29), the density of stained particles (**control** = 0.88 ± 0.18 particles/μm^2^, n= 32 vs **+Ab anti-GluN2B** = 0.97 ± 0.18 particles/μm^2^, n=29), the particle size (**control**=73,387±17,808 nm^2^, n= 32 vs **+Ab anti-GluN2B**=80,299±29,522 nm^2^, n=29), and particle mean fluorescence intensity (**control**=18.19±0.65 A.U., n= 32 vs **+Ab anti-GluN2B**=18.37±1.16 A.U., n=29)(**Fig. 8A**).

**Figure 8.**
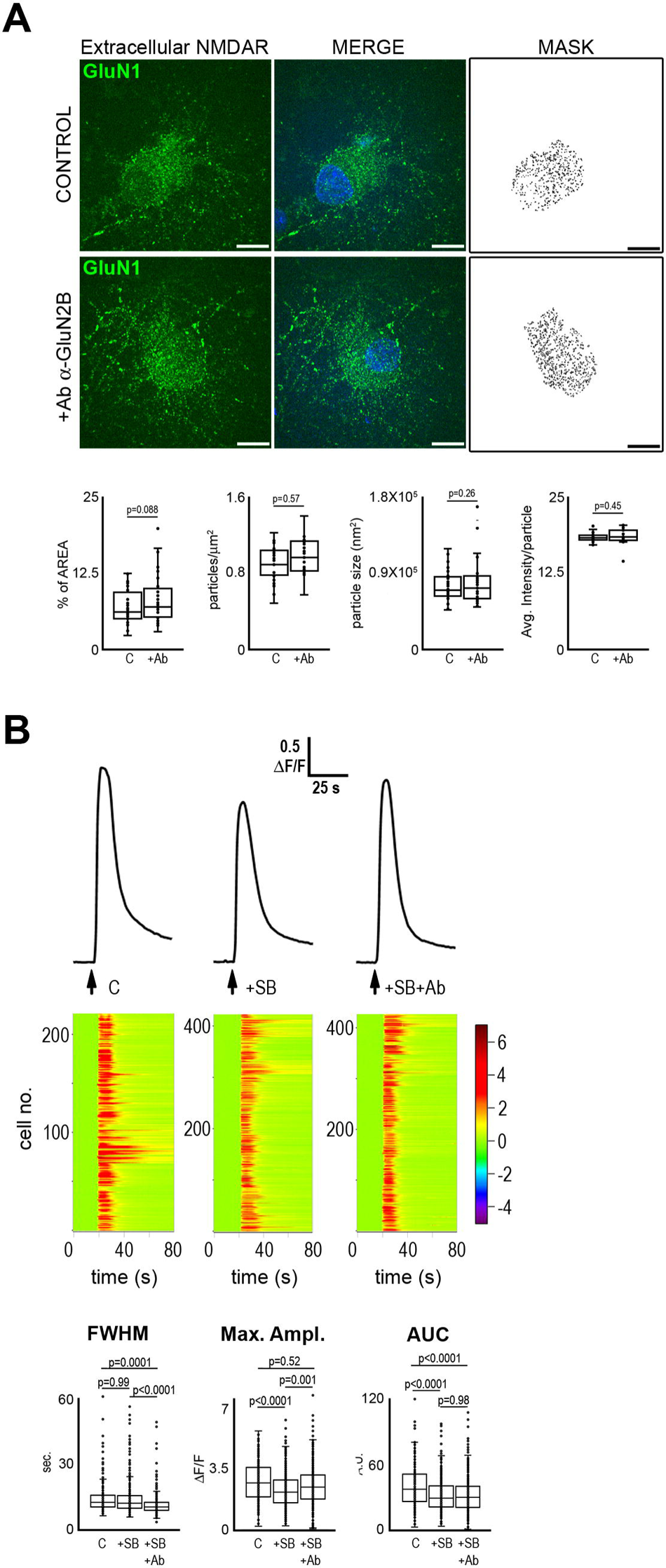
Anti-GluN2B Ab does not reduce surface NMDAR and p38 inhibition prevents Ab effect. *A* Surface NMDAR quantification in control and anti-GluN2B Ab neurons treated 60 min. Merged representative images with stained nuclei and their masks obtained from segmentation are shown in the upper panel. The whisker plots for both conditions for percentage of cell area labeled, particle density, particle size, and averaged fluorescence intensity per particle are shown below. ***B*** Neuronal averaged response to 1 sec NMDA/Gly stimulation (represented by the arrow below the trace) in the control condition (left trace), control condition plus 10 min p38 inhibitor (SB; middle trace), or after 60 min exposure to 5μg/ml control Ab plus SB (right trace). Below each trace, the raster plot of individual cell responses is shown with the same time scale in the *x* axis and the cell number in the *y* axis. The whisker plots for the FWHM, Maximum Amplitude and integrated i[Ca^2+^] (AUC) are shown at the bottom. *p* Student’s t-test values for each comparison are shown above each plot. Reference bar= 20 μm.

These results indicate that NMDAR endocytosis does not occur and are consistent with earlier reports of negligible NMDAR endocytosis in the short-term (see Discussion). Therefore, the reduction of NMDAR-mediated [Ca^2+^]i response induced by the anti-GluN2B Ab after 60 min incubation seems to be unrelated to a diminished density of functional NMDAR at the cell surface.

Then, we tested the role of p38 activation in reducing the NMDAR-mediated [Ca^2+^]i response induced by the anti-GluN2B Ab. Therefore, we tested the effect of the p38 inhibitor SB203580 (SB) on the NMDAR-mediated [Ca^2+^]i response after 60 min anti-GluN2B Ab incubation. As shown in **Fig. 8B**, SB partially prevented the **peak amplitude** reduction (**control** = 2.76 ± 1.17 ΔF/F A.U. vs. **control + SB** = 2.28 ± 1.03 ΔF/F A.U. vs. **a-GluN2B+SB** = 2.55 ± 1.14 A.U.). Nevertheless, the **FWHM** was decreased by the combination of SB and Ab (**control** =15.48 ±7.9 sec. vs. **control+SB** =15.52 ± 12.62 ΔF/F sec. vs. **a-GluN2B+SB** = 12.2 ± 6.72 sec.), resulting in a reduced amount of [Ca^2+^]i (**AUC; control**=39.24±18.95 A.U. vs. **control + SB**=31.68±15.38 A.U. vs. **a-GluN2B+SB** = 31.52 ± 15.96 A.U.).

Notably, SB alone reduced the [Ca^2+^]i response by decreasing the peak amplitude, implicating that basal activity of p38 modulates NMDAR function (see Discussion). These results indicate that p38 activity is involved in the reduction of NMDAR-mediated [Ca^2+^]i response induced by the anti-GluN2B Ab, through regulation of the mean peak amplitude, but also that additional mechanisms are involved.

### Flux-independent NMDAR signaling is involved in anti-GluN2B Ab regulation of NMDAR function

Our experiments so far demonstrated the role of p38 activation in regulating NMDAR function. However, given the different magnitudes of p38 activation by anti-NMDAR Abs, their distinct effects on NMDAR-mediated [Ca^2+^]i response after 60 min and 24 h treatment, and the partial effect of SB, it is also clear that other mechanisms besides p38 activation are involved. Thus, we explored whether NMDAR signaling through Ca^2+^ permeation or flux-independent signaling are involved in the anti-GluN2B Ab effect. Flux-independent signaling has been reported to occur after agonist or co-agonist binding (Montes de Oca-B, 2018).

We first tested whether the anti-GluN2B Ab could induce *per se* channel opening of the NMDAR. While Abs obtained from a-NMDARe patientś CSF did not show this gain of function (Gleichman *et al*., 2012), this mechanism has been demonstrated for Abs against GluN2B in Systemic Lupus Erythematosus (SLE) and other channel-directed Abs involved in autoimmune diseases (Martinez-Martinez et al., 2013). We first recorded cells under basal conditions for 210 sec. Then, the anti-GluN2B Ab was added, and cells were recorded for 40 min. This experiment was performed without perfusion to avoid Ab washout and in culture media to mimic the experimental setting used, in which Abs are incubated in the culture well. As shown in **Fig. 9A**, the anti-GluN2B Ab did not modify baseline [Ca^2+^]i of neurons throughout the recording time. The same was observed in the experiment with RH instead of culture media (**Supporting Fig. 2**). These results demonstrate that the anti-GluN2B Ab does not function as an NMDAR agonist, because a [Ca^2+^]i rise should be observable in all NMDAR of the cell given he high Ab concentration, including those somatic. Therefore, these experiments rule out that the effect of this Ab on NMDAR function is related to channel opening by Ab binding.

**Figure 9.**
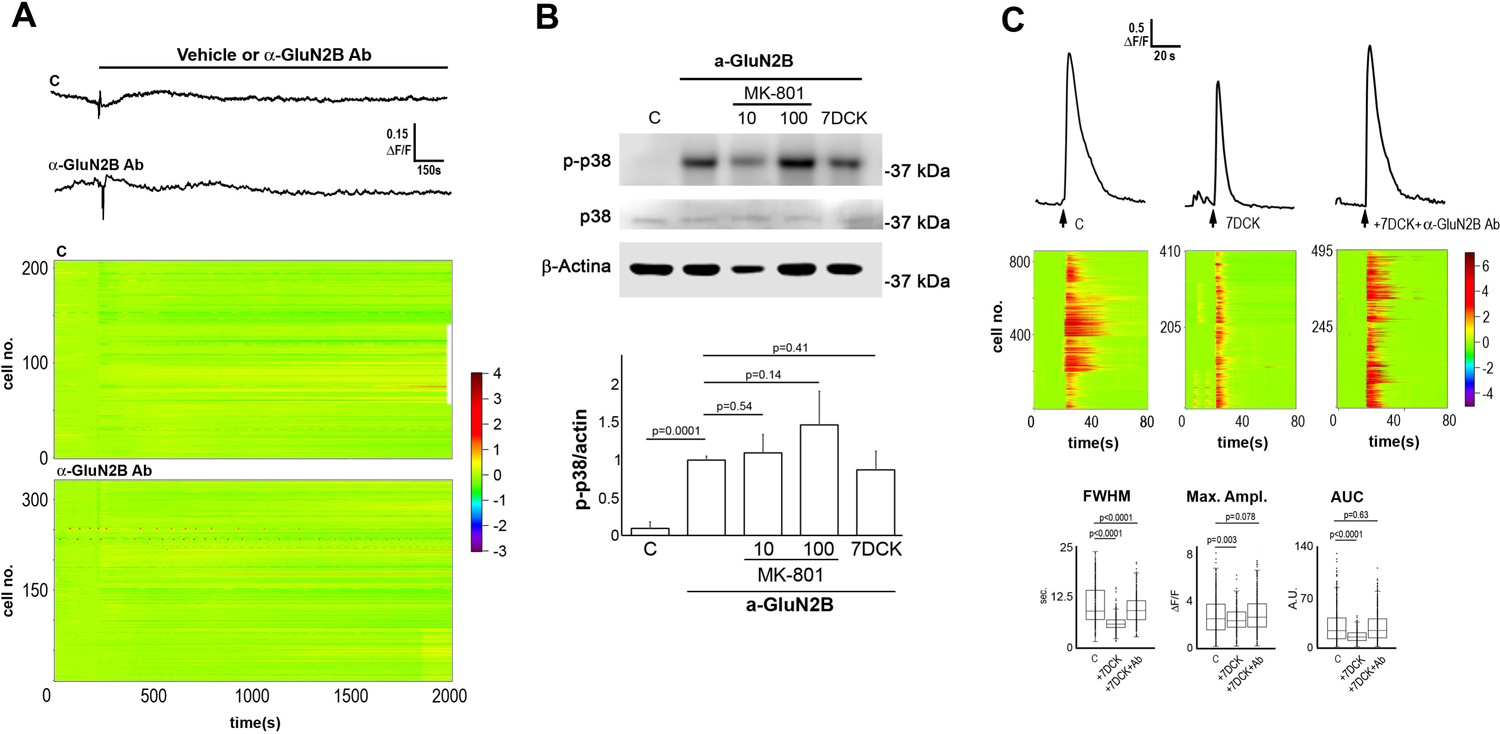
Flux-independent signaling through p38 mediates a-GluN2B reduction of NMDAR-mediated i[Ca^2+^] response. *A* Neuronal i[Ca^2+^] response recorded for 40 min in control conditions (upper trace) or with 5μg/ml anti-GluN2B Ab alone (lower trace) applied after 210 sec basal recording and present throughout the recording (black line above trace). Below the traces, the raster plots of individual cell responses are shown with the same time scale in the *x* axis and the cell number in the *y* axis. ***B*** Western Blot experiment of neuronal lysates treated for 60 min with anti-GluN2B Ab in the presence of NMDAR inhibitors (MK-801 10 μM or 100 μM; 7DCK 100 μM) or control cells. Total p38 and the loading control (actin) are shown in the panels below for comparison. The histogram below shows the average of three independent experiments in which p-p38 band size was quantified and normalized to is loading control. ***C*** Neuronal averaged response to 1 sec NMDA/Gly stimulation (represented by the arrow below the trace) in the control condition (left trace), control condition plus 7DCK (middle trace), or after exposure to 5μg/ml anti-GluN2B Ab plus 7DCK (right trace). Below each trace, the raster plot of individual cell responses is shown with the same time scale in the *x* axis and the cell number in the *y* axis. The whisker plots for the FWHM, Maximum Amplitude and integrated i[Ca2+] (AUC) are shown at the bottom. *p* Student’s t-test values for each comparison are shown above each plot.

Then, we tested the effect of NMDAR inhibitors on the induction of p-p38. We found that NMDAR receptor pore blocker MK-801 (10 μM or 100 μM) did not reduce the amount of p-p38 induced by the anti-GluN2B Ab (**Fig. 9B**). Similarly, the GluN1 specific inhibitor 7DCK (100 μM), that prevents channel opening as well as GluN1 flux-independent signaling (Montes de Oca-B, 2018), neither prevented p-p38 induction (**Fig. 9B**). These findings further support a role of a third-party molecule other than the NMDARs in p38 activation, as suggested by our experiments described above.

Then, we tested the effect of the GluN1 specific inhibitor 7DCK on the down-regulation of NMDAR-mediated [Ca^2+^]i response by anti-GluN2B Ab. Unfortunately, the same experiment with MK-801 was technically unfeasible since it is an irreversible pore blocker. In these experiments we found that 7DCK substantially prevented the reduction of the mean peak amplitude whereas the FWHM was slightly decreased (***FWHM***: **control**=10.67±4.66 sec. vs **7DCK**=6.06±1.94 sec. vs **Ab+7DCK**=9.56±3.22; ***Max. Ampl***.: **control**=2.48±1.37 ΔF/F A.U. vs. **7DCK**=2.21±0.96 vs **Ab+7DCK**=2.64±1.34; ΔF/F A.U.) (**Fig.9C**). This resulted in the prevention of the NMDAR-mediated [Ca^2+^]i reduction by the anti-GluN2B Ab (***AUC*: control**=29.62±23.27 vs. **7DCK**=15.25±7.77 vs **Ab+7DCK**=28.55±20.04). These observations suggest that flux-independent signaling by the NMDAR participates in the regulatory actions of the anti-GluN2B Ab on NMDAR function, since there is no channel opening by this Ab.

## DISCUSSION AND CONCLUSIONS

The a-NMDARe is a recently described autoimmune pathology under intense investigation to elucidate its clinical, cellular, and molecular mechanisms. Although anti-NMDAR Abs are the causing agent of this disease, anti-NMDAR Abs can also be found in patients with psychosis, diabetes, hypertension, SLE, and even in healthy individuals (Hammer *et al*., 2014; Castillo-Gomez *et al*., 2016; Jézéquel *et al*., 2018). Indeed, the occurrence of these Abs in healthy individuals is of great relevance because it has been reported that a-NMDARe could require a compromised blood-brain barrier or meninges to manifest (Diamond *et al*., 2009; Suzuki *et al*., 2013; Hammer *et al*., 2014). These observations led us to investigate the early cellular mechanisms that mediate the effects of anti-NMDAR Abs, which appear to recognize different NMDAR subunits and subunit domains in healthy and diseased seropositive individuals (Castillo-Gomez *et al*., 2016). The elucidation of the mechanisms that these Abs regulate is critical for a deeper understanding of these pathologies and could lead to novel therapeutic approaches.

In this work, we performed our experiments with cultured rat cortical neurons, extensively used to investigate NMDAR function and in which the NMDAR configuration is known. We found that in our model the [Ca^2+^]i response to a brief (1 s) pulse of NMDA(50 μM)/Gly(10 μM) is mainly mediated by the NMDAR, assembled as others have reported (Li *et al*., 1998).

Our experiments showed that Abs targeting the GluN1 IC domain and GluN2B mostly down-regulate NMDAR function or its components after 24 h, as previously demonstrated for a-NMDARe patientś Abs. They also down-regulated NMDAR function after only 60 min of treatment. Only the Ab raised against the extracellular domain of GluN1 slightly increased the amount of [Ca2+]i permeated by the NMDAR at 60 min. In contrast, after 24 h it did not change due to an increase in the mean peak amplitude compensated by a decrease of FWHM (discussed below). Importantly, all these Ab were raised in rabbits, thus reducing the confounding factors due to cross-species interactions. Interestingly, our results demonstrated different outcomes in different assays, depending on the subunit-specific Ab employed, and the duration of the treatment (60 min or 24 h), together indicating that different mechanisms are involved and determine the outcome. Importantly, these observations are unrelated to vehicle or preservatives of the anti-NMDAR Ab because their effect over NMDAR-mediated [Ca^2+^]i or p-p38 induction is not consistent with the volume of Ab or Sodium Azide added to neuronal cultures during treatment (Table 1).

**Table I.** Antibody vial contents according to manufacturer.

| ANTIBODY | CONCENTRATION<br>ug/ml | VEHICLE FROM<br>MANUFACTURER | Sodium Azide % | BSA % | Gelatin % | ul Ab for 300 ul | Final Na Azide % |
| --- | --- | --- | --- | --- | --- | --- | --- |
| anti-GluN1 EC | 1000 | PBS | 0.02 | 1 | 0 | 1.5 | 0.0001 |
| anti-GluN2B EC | 250 | PBS | 0.10 | 0 | 0 | 6 | 0.002 |
| anti-GluN1 IC | 500 | PBS | 0.09 | 0 | 0 | 3 | 0.0009 |
| anti-TrkA | 200 | PBS | 0.10 | 0 | 0.1 | 7 | 0.0023 |

It has been suggested that anti-NMDAR Abs from a-NMDARe patients modulate NMDAR function through steric interactions, regulating the open time of the receptor (Gleichman *et al*., 2012). To test whether this mechanism is involved in our findings, we evaluated whether anti-NMDAR Abs remained bound to the NMDAR of neurons after treatment, labeling and recording. To our surprise, our findings indicate that the steric interaction between the Abs and the NMDAR seems non-relevant in our experimental model, suggesting that other regulatory mechanisms, such as intracellular signaling, may be involved. In addition, the anti-GluN1 IC Ab neither remained attached to the cells, consistently with the intracellular location of its antigen. This Ab neither accessed the cell, as it occurs with most Abs, although a small group of Abs have been reported to do so (Rhodes and Isenberg, 2017; Slastnikova *et al*., 2018). Nevertheless, such cell-penetrating Abs are mostly autoantibodies against DNA or enter the cell bound to antigens of infectious agents. Even though we did not test whether the control Ab employed in Fig. 7 is capable of accessing the cell, given that it does not have the features of those penetrating Abs, we assume that it cannot enter the cell as it is true for most Abs (Rhodes and Isenberg, 2017; Slastnikova *et al*., 2018), and as we have observed in cultured astrocytes (our unpublished observations).

Since steric interactions are not involved, we investigated whether intracellular signaling plays a role. Consistently with this idea, anti-NMDAR Abs induced p-p38 after 60 min treatment. However, strikingly, we also found that an unrelated, non-anti-NMDAR Ab (Control Ab) was also capable of inducing p-p38 and modulating NMDAR function after 60 min treatment. This finding demonstrates that different Abs activate the p38 pathway in our neuronal culture system. Although the p38 pathway is activated by NMDAR flux-independent and -dependent signaling, these results indicate that it can be activated by Abs, probably through an unknown third-party neuronal molecular mediator, because NMDAR inhibitors did not prevent p-p38 induction. Further research is required to elucidate the nature of these putative mediators. Interestingly, FcγRII, receptors for the constant region of Abs, are expressed by different neurons, including cultured rat and mouse cortical neurons (DIV 7) (Kam *et al*., 2013; Fuller, Stavenhagen and Teeling, 2014). Therefore, it is possible that Abs activate FcγRII and indirectly regulate NMDAR function. Although the main intracellular pathways of FcγR are mediated by PLC, PI3K, or Vav/Rho (Fuller, Stavenhagen and Teeling, 2014; Gómez Romána, Murray and Weiner, 2014), p38 has been involved in the intracellular signaling of FcγRII in macrophages (Rose *et al*., 1997; Lucas *et al*., 2005). This possibility requires experimental confirmation and should consider the species of Abs and experimental model employed. If it is the case that FcR mediate Ab regulation of NMDAR function, then its role should be considered in experimental models with cells, living tissues, or live animals, and when cross-species Abs are employed. More importantly, the role of FcR and immune functions of neuroglia could also play a relevant role in Ab-mediated pathologies. These possibilities have not been considered to our knowledge for a-NMDARe or other pathologies and could be critical. Furthermore, since the NMDAR is expressed by neuroglial cells (Hogan-Cann and Anderson, 2016), that in turn modulate neuronal and synaptic function, the effect of anti-NMDAR Abs on these cells should also be investigated. In addition, these findings may shed some light regarding the intriguing observation of pathogenic Abs directed against intracellular molecules in different autoimmune encephalitis (Dalmau and Graus, 2019). Moreover, despite Abs directed against NMDAR intracellular domains have not been described in a-NMDARe, they have been found in other pathologies (Castillo-Gomez *et al*., 2016).

As explained, the chronic a-NMDARe symptoms have been linked to NMDAR hypofunction resulting from the induction of its endocytosis by patient’s Abs. Nevertheless, the early mechanisms involved have not been fully established. This is critical since the early diagnosis could condition disease phenotype and severity and could provide new opportunities for effective therapies. Given that contradictory findings have been reported regarding the endocytosis of the NMDAR in response to <3 h treatment with a-NMDARe patientś Abs, we investigated if 60 min treatment with anti-GluN2B Ab decreased surface NMDAR in our culture model. We did not detect a change of surface NMDAR with all parameters evaluated, only a trend to augment the percentage of the area was observed, similar to a previous report where larger and more abundant NMDAR clusters were found by STORM microscopy (Ladepeche *et al*., 2018). Thus, no endocytosis is induced by anti-GluN2B Ab in our model in this short time, consistently with previous reports suggesting that NMDAR endocytosis can be detected by similar methods only after >12 h incubation with a-NMDARe Abs.

Our data with the anti-GluN2B Ab also demonstrated that p38 activation has a role regulating NMDAR function. In particular, the time component, since SB, a p38 inhibitor, prevented its reduction induced by the Ab. Despite all anti-NMDAR and control Abs elicited p-p38, the level of induction varied among them. The Abs directed against the extracellular domain of the NMDAR induced moderate p-p38, whereas the pair of Abs that could not access their intended target induced massive p-p38. These results suggests that other mechanisms are involved in the modulation of the p38 pathway, besides the putative third-party mediator. In this regard, Ab binding to the extracellular domain of the NMDAR and its oligomerization during treatment may play a role limiting the amount of p-p38. However, this needs to be tested. Interestingly, the reduction of NMDAR function by SB alone indicates that basal p38 activity regulates it, an observation not previously reported to the extent of our knowledge, thus forming a regulatory loop, since NMDAR induces p-p38 (Waxman and Lynch, 2005). p38 is a serine/threonine kinase that could phosphorylate the NMDAR at serine and threonine residues known to regulate its function (Traynelis *et al*., 2010).

In addition, our experiments showed that the anti-GluN2B Ab did not work as an NMDAR agonist, as it has been found for other receptor-targeted Abs with a gain of function (Martinez-martinez *et al*., 2013). We also found that NMDAR inhibitors that block ion flux (MK-801) or channel opening through co-agonist binding inhibition (7DCK), did not prevent p-p38 induction. These observations further support the notion that a third-party mediator is responsible for p-p38 activation. Even though 7DCK did not reduce p-p38, it prevented the reduction of the NMDAR-mediated [Ca^2+^]i response induced by the anti-GluN2B Ab. These findings suggest that Ab binding to the NMDAR stimulates GluN1 flux-independent signaling, because no [Ca^2+^] entry was promoted by anti-GluN2B Ab, but still 7DCK prevented its effect. Also, they indicate that in our experimental conditions the p38 pathway and flux-independent signaling synergize for NMDAR regulation. The role of NMDAR flux-independent signaling is further indicated by the reduction of the NMDAR-mediated [Ca^2+^]i response by 7DCK alone. This finding indicates that GluN1 flux-independent signaling can regulate the NMDAR function itself, implicating a regulatory loop that had not been described. It is important to note that in our experimental design (with or without anti-GluN2B, although see below), no agonist for the opening of the NMDAR is present in culture media during Ab treatment. Therefore, it can be assumed that the specificity of the response elicited by this anti-GluN2B Ab compared with the other Abs employed, could result also from NMDAR-Ab interaction and NMDAR oligomerization.

Flux-independent signaling by the NMDAR signaling was described almost 20 years ago for its ability to induce NMDAR endocytosis (Vissel *et al*., 2001; Nong *et al*., 2003). This pathway is getting significant attention as it may be involved in physiological and pathological processes such as LTD, Alzheimeŕs Disease or schizophrenia (Montes de Oca-B, 2018, 2022; Park, Stein and Zito, 2022). In fact, flux-independent signaling has also been reported for other ionotropic receptors as mentioned above, and it has been suggested that it could be a common mechanism elicited by ionotropic receptors. Importantly, flux-independent NMDAR signaling by the NMDAR has been shown to mediate other mechanisms besides NMDAR endocytosis, such as LTD, synaptic depression induced by beta-amyloid, Ca^2+^ release from intracellular pools, and plasma membrane-mitochondria interactions (Reviewed in Montes de Oca-B, 2018; Montes de Oca Balderas et al., 2020). Thus, it is possible that this signaling mechanism could be profited for therapeutics of a-NMDARe patients as it has been shown for cerebral ischemia in animal models (Chen *et al*., 2017).

A relevant question is how anti-GluN2B Ab elicits flux-independent signaling by the NMDAR. We currently envisage two possibilities. The first one is that the anti-GluN2B Ab induces a conformational change of GluN2B, that could itself generate flux-independent signaling that requires to be investigated, that does not open the channel but transfers conformational information to GluN1 that elicits flux-independent signaling (Dore, Aow and Malinow, 2015; Hansen *et al*., 2018; Montes de Oca-B, 2018). The second non-mutually exclusive possibility is that Gly present in the Neurobasal culture media may induce such signaling through GluN1binding. Therefore, when 7DCK alone is added, such signaling is blocked and the NMDAR function is down-regulated. However, this putative pathway has a different outcome when the Ab is present. Nonetheless, it is still possible to explain these observations without the participation of flux-independent signaling. Spontaneous synaptic neurotransmitter release may occur that elicit NMDAR activation and an increase of local [Ca^2+^]i. If 7DCK blocks such flux-dependent synaptic events, it could cause NMDAR down-regulation. However, there are two concerns regarding this possibility. First, it must be noted that with our methodology we measure the [Ca^2+^]i signal from the soma of the cell, response that is mainly given by the NMDA/Gly challenge of extrasynaptic, mainly somatic NMDAR. Then, this possibility implicates that the few small, spontaneous, local, synaptic events (See Fig. 9A) regulate the function of synaptic, extrasynaptic and somatic NMDAR. Second, if the down-regulation of synaptic events leads to down-regulation of extrasynaptic and somatic NMDAR, it would implicate a negative feedback loop. However, this loop was not observed in presence of the anti-GluN2B Ab and 7DCK. Therefore, an effect of 7DCK on all NMDAR blocking their flux-independent signaling is favored instead of a specific, constrained effect restricted to synaptic NMDAR. However, further research is required to confirm if this mechanism of anti-NMDAR Ab action occurs with a-NMDARe patientś Abs.

Interestingly, our results also suggest opposing actions of p38 in NMDAR regulation. On the one hand, basal p38 activity is required for somatic NMDAR function, the population mainly activated in response to NMDA/Gly in our experimental conditions. On the other hand, during the anti-GluN2B Ab challenge, p38 mediates NMDAR down-regulation. These opposing actions could be related to p38 autophosphorylation and/or the two mechanisms of p38 activation: through the classical phosphorylation at Thr^180^/Tyr^182^ assayed here, or the alternative pathway through phosphorylation of Tyr^323^ by Lck and ZAP70 cascades described in T cells (Salvador *et al*., 2005), mechanisms that have opposing intracellular effects (Alam *et al*., 2014). This double-sided effect of p38 inhibition indicates that it plays a homeostatic role also for NMDAR function. Regarding this double-sided function of p38 in neurons, the role of astrocytes and their peri-synaptic projections could be relevant because they have been shown to orchestrate neuronal function and to secrete D-Ser, co-agonist of the NMDAR necessary for LTP, that requires Ca^2+^ permeation, whereas flux-independent signaling can be elicited with the co-agonist or the agonist alone (Halassa and Haydon, 2010; Henneberger *et al*., 2010; De Pitta, Brunel and Volterra, 2016; Montes de Oca-B, 2018).

Different groups have tackled the early effects of anti-NMDAR Abs with distinct experimental approaches, but as explained above, contradictory findings have been reported. This lack of consensus could be linked with different causes including the cell or tissue model employed, the expression of putative third-party mediators in such models (i.e. FcR, as shown here), the methodological approaches, the NMDAR subunit configuration employed in these approaches, Ab steric competition, the use of different Ab sources such as CSF or isolated IgG from serum or CSF, different Ig isotypes and/or affinities, or interindividual differences. Also, the diversity and relative abundance of IgG clones directed against different subunits, domains, and epitopes of NMDAR subunits in patientś fluids could also contribute as a source of variability, as suggested by our experiments. Some studies have suggested that anti-GluN1 Abs are responsible for the a-NMDARe phenotype. However, the presence of anti-GluN2B Ab has not been entirely ruled out since some factors have been disregarded (Dalmau *et al*., 2008; Gleichman *et al*., 2012; Castillo-Gomez *et al*., 2016; Malviya *et al*., 2017). These factors may include information transfer among NMDAR subunits (Dore, Aow and Malinow, 2015; Hansen *et al*., 2018; Montes de Oca-B, 2018), subunit-specific conformational rearrangements in intracellular compartments or at the cell surface after heteromeric association in the Endoplasmic Reticulum, or specific conformations with single subunit homomeric transfection.

Moreover, one must bear in mind that endogenous NMDAR are assembled with two GluN1 and GluN2 subunits (Hansen *et al*., 2018), that ovarian teratomas express different NMDAR subunits (Tachibana *et al*., 2010; Tabata *et al*., 2014), that human population presents a diversity of immune reactions that may result into a diversity of Abs and Abs specificities as has been suggested (Malviya *et al*., 2017), that an anti-GluN1 monoclonal Ab reconstructed from a patient does not fully compete for antigenic sites recognized by Ab in patientś CSF (Malviya *et al*., 2017), and that Abs against GluN1/GluN2B heteromers were initially described in patients of this pathology (Dalmau *et al*., 2007; Iizuka *et al*., 2008; Tachibana *et al*., 2010; Tabata *et al*., 2014). Furthermore, it is intriguing how the immune system can discriminate different NMDAR subunits and develop only a single GluN1 targeted immune reaction while disregarding the other subunits. In this regard, up to 7 different peptides from the extracellular domain of GluN1 have been found as major immunogenic determinants, three of which contain critical residues for Gly binding (During *et al*., 2000). Thus, it seems challenging that a single antigenic site is responsible for the disease phenotype, rather than the induction of anti-NMDAR Ab populations that affect the entire NMDAR structure-function. Taken together, our findings and these considerations strongly suggest that subpopulations of anti-GluN1 and other subunits may exist in the pool of Abs of a-NMDARe patients, as suggested by others (Malviya *et al*., 2017), that nevertheless may be challenging to evidence.

Oddly, as mentioned above, the anti-GluN1 Ab generated a unique response pattern. At 24 h it decreased FWHM but augmented the peak amplitude resulting in no net gain or loss of [Ca^2+^]i. At 60 min it increased FWHM but decreased peak amplitude, resulting in the increase of [Ca^2+^]i. In contrast, all other Abs tested decreased the [Ca^2+^]i response in all conditions. This response by the anti-GluN1 was unexpected because it has been shown that anti-GluN1 Abs induce NMDAR endocytosis, and it has been suggested that all-natural occurring anti-GluN1 Abs have pathogenic potential (Castillo-Gomez *et al*., 2016).

Nevertheless, our findings suggest that this anti-GluN1 Ab may also alter NMDAR and cell function given the regulation of time and peak components, without affecting the total amount of mobilized Ca^2+^. This is because spike and time amplitudes determine intracellular signaling and their modification may implicate a distinct cellular outcome. These findings indicate that subunit-specific Abs may modify NMDAR function by different mechanisms. It is worth noticing that this anti-GluN1 Ab is raised against a sequence of 100 N-terminal aa out of 540 aa that comprises the EC domain. Thus, as others have shown, the region recognized by the Ab impacts the outcome. In this regard, two key aa (N^368^/G^369^) (glycosylation site 7; G7) of GluN1 have been identified as critical for the binding of patientś anti-GluN1 Abs (Gleichman *et al*., 2012; Castillo-Gomez *et al*., 2016). These observations could be relevant for anti-NMDAR seropositive healthy individuals and patients with other pathologies. Although the mechanism behind anti-GluN1 Ab modulation of NMDAR function was not investigated, according to our results, the steric modulation of the open probability of the NMDAR, that has been reported previously (Gleichman *et al*., 2012), does not occur since the Ab does not remain bound after cell treatment, labeling and recording. However, modulation of the open probability may also stem from NMDAR post-translational modifications resulting from intracellular transduction pathways (Traynelis *et al*., 2010).

All in all, our results demonstrate that anti-NMDAR Abs can regulate NMDAR function after <3h incubation through different mechanisms, including intracellular signaling. More importantly, our results suggest that an unknown, third-party mediator expressed by neurons is involved in the activation of p38 by anti-NMDAR and non-anti-NMDAR Abs. These findings are relevant mainly considering the cerebral setting, where neuroglia and the Blood Brain Barrier (BBB) are critical for the immune response. Interestingly, our results also suggest that flux-independent NMDAR signaling is involved in NMDAR regulation by anti-GluN2B Ab and could be responsible for its endocytosis observed in the long-term. Finally, our findings with subunit-specific Ab could help to set standards for better a-NMDARe diagnosis, since certain symptoms may be related to Ab targeting of specific subunits or NMDAR subunit configurations. Also, they help to understand this disease better, and possibly in the future, to ideate novel therapies. However, further research is required to confirm if these mechanisms of anti-NMDAR Abs occur with a-NMDARe patientś Abs.

## DATA AVAILABILITY

All data described in this manuscript are contained within the manuscript.

## ACKNOWLWEDGEMENTS

PMOB deeply thanks his family that has supported him despite lack of conditions to do science at INNN and in general in México. Authors thank Teresa Montiel, Beatriz Aguilar Maldonado, and Dr. Nicolas Jiménez Pérez for their technical assistance; Claudia Rivera and Héctor Malagón for animal facilities; Francisco Pérez Eugenio from the Computing Unit of IFC; and Dr. Teresa Romero and Dr. Norma Sánchez for helping us with Abs or reagents. PMOB thanks Dr. Guillermo Hernández Mendoza for his help to develop the MATHLAB code. This work received funding from DGAPA. No. IG200119; CONACYT No. 315803; CONACYT No. 21887 to AHC and CONACYT No. 132706 to PMOB. Also, from grant A1S-17357 and PAPIIT-UNAM grant IN204919 to LMT. JCGG is a Master student from the Programa de Maestría y Doctorado en Ciencias Bioquímicas, at the Universidad Nacional Autónoma de México (UNAM) and he was a recipient of CONACyT fellowship (CVU 852732).

---

**Supplementary Figure 1.** anti-NMDAR Ab induce p38 phosphorylation and a ∼53 kDa band detected with the monoclonal anti-phospho-p38 Ab. Western Blot experiment of neuronal lysates treated for 60 min with the three anti-NMDAR Ab or control cells. The loading control (actin) is shown in the panel below for comparison. One representative experiment is shown of at least three independent performed.

**Supplementary Figure 2.** Anti-GluN2B Ab does not induce i[Ca2+] response in RH. Neuronal i[Ca^2+^] response to 5μg/ml a-GluN2B Ab alone applied during 10 sec (arrow) and present throughout the recording in RH. The arrowhead shows the application of a high KCl (300 mM), to test neuronal responsiveness. Below the trace, the raster plot of individual cell responses is shown with the same time scale in the *x* axis and the cell number in the *y* axis.

## Notes

### Competing Interest Statement

The authors have declared no competing interest.

## REFERENCES

Alam, M. S. et al. (2014) ‘Counter-regulation of T cell effector function by differentially activated p38’, Journal of Experimental Medicine, 211(6), pp. 1257–1270. doi: 10.1084/jem.20131917.

Amrhein, V., Greenland, S. and Mcshane, B. (2019) ‘Scientists rise up against statistical significance’, Nature, 567(21 march), pp. 305–307.

Atlas, D. (2014) ‘Voltage-gated calcium channels function as Ca-activated signaling receptors’, Trends in Biochemical Sciences. Elsevier Ltd, 39(2), pp. 45–52. doi: 10.1016/j.tibs.2013.12.005.

Atlas, D. (2022) ‘Revisiting the molecular basis of synaptic transmission’, Progress in Neurobiology. Elsevier Ltd, 216(June), p. 102312. doi: 10.1016/j.pneurobio.2022.102312.

Castillo-Gomez, E. et al. (2016) ‘All naturally occurring autoantibodies against the NMDA receptor subunit NR1 have pathogenic potential irrespective of epitope and immunoglobulin class’, Molecular Psychiatry. Nature Publishing Group, 22(12), pp. 1776– 1784. doi: 10.1038/mp.2016.125.

Chen, J. et al. (2017) ‘A non-ionotropic activity of NMDA receptors contributes to glycine-induced neuroprotection in cerebral ischemia-reperfusion injury’, Scientific Reports. Springer US, 7(January), pp. 1–14. doi: 10.1038/s41598-017-03909-0.

Chung, C. (2013) ‘NMDA receptor as a newly identified member of the metabotropic glutamate receptor family: Clinical implications for neurodegenerative diseases’, Molecules and Cells, 36(2), pp. 99–104. doi: 10.1007/s10059-013-0113-y.

Cidad, P. et al. (2012) ‘Kv1.3 Channels Can Modulate Cell Proliferation during Phenotypic switch by an ion-Flux independent Mechanism’, *Arterioscler Thromb Vasc Biol*, May, pp. 1299–1307. doi: 10.1161/ATVBAHA.111.242727.

Dai, J. et al. (2021) ‘GluD1 is a signal transduction device disguised as an ionotropic receptor’, Nature. Springer US, 595(July), pp. 261–265. doi: 10.1038/s41586-021-03661-6.

Dalmau, J. et al. (2007) ‘Paraneoplastic Anti – N-methyl-D-aspartate Receptor Encephalitis Associated with Ovarian Teratoma’, Annals of Neurology, 61(1), pp. 25–36. doi: 10.1002/ana.21050.

Dalmau, J. et al. (2008) ‘Anti-NMDA-receptor encephalitisL: case series and analysis of the eff ects of antibodies’, The Lancet Neurology, 7(Dec), pp. 1091–1098. doi: 10.1016/S1474-4422(08)70224-2.

Dalmau, J. et al. (2011) ‘Clinical experience and laboratory investigations in patients with anti-NMDAR encephalitis’, The Lancet Neurology. Elsevier Ltd, 10(1), pp. 63–74. doi: 10.1016/S1474-4422(10)70253-2.

Dalmau, J. and Graus, F. (2019) ‘Antibody-Mediated Encephalitis’, The New England Journal of Medicine, 378(9), pp. 840–851. doi: 10.1056/NEJMra1708712.

Diamond, B. et al. (2009) ‘Losing your nerves? Maybe it’s the antibodies’, Nature Reviews Immunology, 9(June), pp. 449–456.

Dore, K. et al. (2017) ‘Unconventional NMDA Receptor Signaling’, The Journal of neuroscience_J: the official journal of the Society for Neuroscience, 37(45), pp. 10800– 10807. doi: 10.1523/JNEUROSCI.1825-17.2017.

Dore, K., Aow, J. and Malinow, R. (2015) ‘Agonist binding to the NMDA receptor drives movement of its cytoplasmic domain without ion flow’, Proceedings of the National Academy of Sciences, 112(47), pp. 14705–14710. doi: 10.1073/pnas.1520023112.

During, M. J. et al. (2000) ‘An Oral Vaccine Against NMDAR1 with Efficacy in Experimental Stroke and Epilepsy’, Science, 287(February), pp. 1453–1461.

Edelstein, A. et al. (2010) ‘Computer Control of Microscopes Using μ Manager’, *Current Protocols Mol*. Biol., 92(1), pp. 14.20.1-14.20.17. doi: 10.1002/0471142727.mb1420s92.

Fernández-Tenorio, M. et al. (2011) ‘Metabotropic regulation of RhoA/Rho-associated kinase by l-type Ca2+ Channels: New Mechanism for Depolarization-Evoked Mammalian Arterial Contraction’, Circulation Research, 108(11), pp. 1348–1357. doi: 10.1161/CIRCRESAHA.111.240127.

Fuller, J., Stavenhagen, J. B. and Teeling, J. L. (2014) ‘New roles for Fc receptors in neurodegeneration-the impact on Immunotherapy for Alzheimer’s Disease’, Frontiers in neuroscience, 8(August), pp. 1–10. doi: 10.3389/fnins.2014.00235.

Gérard, F. and Hansson, E. (2012) ‘Inflammatory activation enhances NMDA-triggered Ca2 signalling and IL-1β secretion in primary cultures of rat astrocytes’, Brain Research, 1473, pp. 1–8. doi: 10.1016/j.brainres.2012.07.032.

Gleichman, A. J. et al. (2012) ‘Anti-NMDA Receptor Encephalitis Antibody Binding Is Dependent on Amino Acid Identity of a Small Region within the GluN1 Amino Terminal Domain’, Journal of Neuroscience, 32(32), pp. 11082–11094. doi: 10.1523/JNEUROSCI.0064-12.2012.

Gómez Romána, V. R., Murray, J. C. and Weiner, L. M. (2014) Antibody-Dependent Cellular Cytotoxicity (ADCC)in ANTIBODY Fc: LINKING ADAPTIVE AND INNATE IMMUNITY. First. Edited by M. E. Ackerman and NimmerjahnFalk. London: Academic Press.

Gómora-García, J. C. et al. (2021) ‘IRE1 α RIDD activity induced under ER stress drives neuronal death by the degradation of 14-3-3 θ mRNA in cortical neurons during glucose deprivation’, Cell Death Discovery. Springer US, 7(131), pp. 1–15. doi: 10.1038/s41420-021-00518-9.

Gray, J. A., Zito, K. and Hell, J. W. (2016) ‘Non-ionotropic signaling by the NMDA receptor: Controversy and opportunity [version 1; referees: 2 approved]’, F1000Research, 5(0), pp. 1–8. doi: 10.12688/F1000RESEARCH.8366.1.

Halassa, M. M. and Haydon, P. G. (2010) ‘Integrated brain circuits: astrocytic networks modulate neuronal activity and behavior’, Annu Rev Physiol, 72, pp. 335–355. Available at: http://www.ncbi.nlm.nih.gov/entrez/query.fcgi?cmd=Retrieve&db=PubMed&dopt=Citation&list_uids=20148679.

Hammer, C. et al. (2014) ‘Neuropsychiatric disease relevance of circulating anti-NMDA receptor autoantibodies depends on blood – brain barrier integrity’, Molecular Psychiatry, 19(September 2013), pp. 1143–1149. doi: 10.1038/mp.2013.110.

Hansen, K. B. et al. (2018) ‘Structure, function, and allosteric modulation of NMDA receptors’, JGP, 150(8), pp. 1081–1105.

Hardingham, G. E. and Bading, H. (2010) ‘Synaptic versus extrasynaptic NMDA receptor signalling: implications for neurodegenerative disorders’, Nat Rev Neurosci, 11(10), pp. 682–696. Available at: http://www.ncbi.nlm.nih.gov/entrez/query.fcgi?cmd=Retrieve&db=PubMed&dopt=Citation&list_uids=20842175.

Henneberger, C. et al. (2010) ‘Long-term potentiation depends on release of D-serine from astrocytes’, Nature, 463(January), pp. 232–236. doi: 10.1038/nature08673.

Hogan-Cann, A. D. and Anderson, C. M. (2016) ‘Physiological Roles of Non-Neuronal NMDA Receptors’, Trends in Pharmacological Sciences. Elsevier Ltd, 37(9), pp. 750–767. doi: 10.1016/j.tips.2016.05.012.

Hughes, E. G. et al. (2010) ‘Cellular and synaptic mechanisms of anti-NMDA receptor encephalitis’, Journal of Neuroscience, 30(17), pp. 5866–5875. doi: 10.1523/JNEUROSCI.0167-10.2010.

Iizuka, T. et al. (2008) ‘Anti-NMDA receptor encephalitis in Japan Long-term outcome without tumor removal’, Neuirology, 70(Feb 12), pp. 504–511.

Jézéquel, J. et al. (2018) ‘Pathogenicity of Antibodies against NMDA Receptor: Molecular Insights into Autoimmune Psychosis’, Trends in Neurosciences, 41(8), pp. 502–511. doi: 10.1016/j.tins.2018.05.002.

Jiménez-pérez, L. et al. (2016) ‘Molecular Determinants of Kv1.3 Potassium Channels-induced Proliferation’, JBC, 291(7), pp. 3569–3580. doi: 10.1074/jbc.M115.678995.

Kaczmarek, L. K. (2006) ‘Non-conducting functions of voltage-gated ion channels’, Nature Reviews Neuroscience, 7, pp. 6–10. doi: 10.1038/nrn1988.

Kam, T. et al. (2013) ‘impairment in Alzheimer’s disease Fc γ RIIb mediates amyloid-β neurotoxicity and memory impairment in Alzheimer’s disease’, Journal of Clinical Investigation, 123(7), pp. 2791–2802. doi: 10.1172/JCI66827.identified.

Kessels, H. W., Nabavi, S. and Malinow, R. (2013) ‘Metabotropic NMDA receptor function is required for beta-amyloid-induced synaptic depression’, Proc Natl Acad Sci U S A, 110(10), pp. 4033–4038. Available at: http://www.ncbi.nlm.nih.gov/entrez/query.fcgi?cmd=Retrieve&db=PubMed&dopt=Citation&list_uids=23431156.

Ladepeche, L. et al. (2018) ‘NMDA Receptor Autoantibodies in Autoimmune Encephalitis Cause a Subunit-Specific Nanoscale Redistribution of NMDA Receptors NMDA Receptor Autoantibodies in Autoimmune Encephalitis Cause a Subunit-Specific Nanoscale Redistribution of NMDA Receptors’, Cell reports, 23(Jun26), pp. 3759–3768. doi: 10.1016/j.celrep.2018.05.096.

Li, J. H. et al. (1998) ‘Developmental changes in localization of NMDA receptor subunits in primary cultures of cortical neurons’, European Journal of Neuroscience, 10(December 1997), pp. 1704–1715.

Lucas, M. et al. (2005) ‘ERK Activation Following Macrophage Fc_R Ligation Leads to Chromatin Modifications at the IL-10 Locus’, Journal of Immunology, 175, pp. 469–477. doi: 10.4049/jimmunol.175.1.469.

Malviya, M. et al. (2017) ‘NMDAR encephalitisL: passive transfer from man to mouse by a recombinant antibody’, Annals of Clinical and Traslational Neurology, 4(11), pp. 768– 783. doi: 10.1002/acn3.444.

Martinez-martinez, P. et al. (2013) ‘Autoantibodies to neurotransmitter receptors and ion channelsL: from neuromuscular to neuropsychiatric disorders’, Frontiers in genetics, 4(September), pp. 1–8. doi: 10.3389/fgene.2013.00181.

Mikasova, L. et al. (2012) ‘Disrupted surface cross-talk between NMDA and Ephrin-B2 receptors in anti-NMDA encephalitis’, Brain, 135(5), pp. 1606–1621. doi: 10.1093/brain/aws092.

Montes de Oca-B, P. (2018) ‘Flux-Independent NMDAR SignalingL: Molecular Mediators, Cellular Functions, and Complexities’, Int J Mol Sci, 19(3800). doi: 10.3390/ijms19123800.

Montes de Oca-B, P. (2022) ‘Meeting reportL: Flux-independent signaling by ionotropic receptorsL: Unforeseen roles, complexities, and challenges’, Journal of Biological Chemistry. The Author, 298(9), p. 102330. doi: 10.1016/j.jbc.2022.102330.

Montes de Oca Balderas, P., et al. (2020) ‘NMDAR in cultured astrocytesL: Flux-independent pH sensor and flux-dependent regulator of mitochondria and plasma membrane-mitochondria bridging’, The FASEB Journal, 34(12), pp. 16622–16644. doi: 10.1096/fj.202001300R.

Montes de Oca Balderas, P. and Aguilera, P. (2015) ‘A Metabotropic-Like Flux-Independent NMDA Receptor Regulates Ca2+ Exit from Endoplasmic Reticulum and Mitochondrial Membrane Potential in Cultured Astrocytes.’, PloS one, 10(5), p. e0126314. doi: 10.1371/journal.pone.0126314.

Montes de Oca Balderas, P. and Gonzalez Hernandez, J. R. (2018) ‘NMDA Receptors in Astroglia: Chronology, Controversies and Contradictions from a Complex Molecule’, in Maria Teresa Gentile and D’Amato, L. C. (eds) Astrocyte Physiology and Pathology. 1st edn. Intech Open.

Moscato, E. H. et al. (2014) ‘Acute mechanisms underlying antibody effects in anti-N-methyl-D-aspartate receptor encephalitis’, Annals of Neurology, 76(1), pp. 108–119. doi: 10.1002/ana.24195.

Nabavi, S. et al. (2013) ‘Metabotropic NMDA receptor function is required for NMDA receptor-dependent long-term depression’, Proc Natl Acad Sci U S A, 110(10), pp. 4027– 4032. Available at: http://www.ncbi.nlm.nih.gov/entrez/query.fcgi?cmd=Retrieve&db=PubMed&dopt=Citation&list_uids=23431133.

Nong, Y. et al. (2003) ‘Glycine binding primes NMDA receptor internalization’, Nature, 422(6929), pp. 302–307. Available at: http://www.ncbi.nlm.nih.gov/entrez/query.fcgi?cmd=Retrieve&db=PubMed&dopt=Citation&list_uids=12646920.

Paoletti, P., Bellone, C. and Zhou, Q. (2013) ‘NMDA receptor subunit diversity: impact on receptor properties, synaptic plasticity and disease’, Nat Rev Neurosci, 14(6), pp. 383–400. Available at: http://www.ncbi.nlm.nih.gov/entrez/query.fcgi?cmd=Retrieve&db=PubMed&dopt=Citation&list_uids=23686171.

Park, D. K., Stein, I. S. and Zito, K. (2022) ‘Ion flux-independent NMDA receptor signaling’, Neuropharmacology. Elsevier Ltd, 210(March), p. 109019. doi: 10.1016/j.neuropharm.2022.109019.

De Pitta, M., Brunel, N. and Volterra, A. (2016) ‘Astrocytes: Orchestrating synaptic plasticity?’, Neuroscience, 323(May 26), pp. 43–61. doi: 10.1016/j.neuroscience.2015.04.001.

Rhodes, D. A. and Isenberg, D. A. (2017) ‘TRIM21 and the Function of Antibodies inside Cells’, Trends in Immunology. Elsevier Ltd, 38(12), pp. 916–926. doi: 10.1016/j.it.2017.07.005.

Richter, K. et al. (2016) ‘Phosphocholine – an agonist of metabotropic but not of ionotropic functions of α 9-containing nicotinic acetylcholine receptors’, Scientific Reports. Nature Publishing Group, 6(28660), pp. 1–13. doi: 10.1038/srep28660.

Rodrigues, R. J. and Lerma, J. (2012) ‘Metabotropic signaling by kainate receptors’, Neuron, 1(August), pp. 399–410. doi: 10.1002/wmts.35.

Rodriguez-Moreno, A. and Lerma, J. (1998) ‘Kainate Receptor Modulation of GABA Release Involves a Metabotropic Function’, Neuron, 20, pp. 1211–1218.

Rose, D. et al. (1997) ‘Fcy Receptor Cross-Linking Activates p42, p38, and JNK/SAPK Mitogen-Activated Protein Kinases in Murine Macrophages’, Journal of Immunology, 158, pp. 3433–3438.

Rubio-Augusti, I. et al. (2011) ‘Isolated hemidistonia associated with NMDA receptor antibodies.’, Mov Dis, 26(2), pp. 351–352.

Salvador, J. M. et al. (2005) ‘Alternative p38 activation pathway mediated by T cell receptor – proximal tyrosine kinases’, Nature immunology, 6(4), pp. 390–395. doi: 10.1038/ni1177.

Schneider, C.A., Rasband, W. S. and Eliceiri, K. W. (2012) ‘NIH Image to ImageJ: 25 years of image analysis.’, Nature Methods, 9, pp. 671–675.

Slastnikova, T. A. et al. (2018) ‘Targeted Intracellular Delivery of AntibodiesL: The State of the Art’, 9(October), pp. 1–21. doi: 10.3389/fphar.2018.01208.

Stein, I. S. et al. (2020) ‘Molecular Mechanisms of Non-ionotropic NMDA Receptor Signaling in Dendritic Spine Shrinkage’, Journal of Neuroscience, 40(19), pp. 3741–3750.

Stein, I. S. et al. (2021) ‘Non-ionotropic NMDAR signaling gates bidirectional structural plasticity in dendritic spines’, Cell reports, 34(January 26), pp. 1–9.

Stein, I. S., Gray, J. A. and Zito, K. (2015) ‘Non-Ionotropic NMDA Receptor Signaling Drives Activity-Induced Dendritic Spine Shrinkage’, Journal of Neuroscience, 35(35), pp. 12303–12308. doi: 10.1523/JNEUROSCI.4289-14.2015.

Suzuki, H. et al. (2013) ‘Anti-NMDAR encephalitis preceded by dura mater lesions’, Neurological Sciences, 34, pp. 1021–1022. doi: 10.1007/s10072-012-1169-8.

Tabata, E. et al. (2014) ‘Immunopathological Significance of Ovarian Teratoma in Patients with Anti-N-Methyl-D-Aspartate Receptor Encephalitis’, European Neurolog, 71, pp. 42–48. doi: 10.1159/000353982.

Tachibana, N. et al. (2010) ‘Expression of Various Glutamate Receptors Including N-Methyl-D-Aspartate Receptor (NMDAR) in an Ovarian Teratoma Removed from a Young Woman with Anti-NMDAR Encephalitis’, Internal Medicine, 49, pp. 2167–2173. doi: 10.2169/internalmedicine.49.4069.

Tamburri, A. et al. (2013) ‘NMDA-receptor activation but not ion flux is required for amyloid-beta induced synaptic depression’, PLoS One, 8(6), p. e65350. Available at: http://www.ncbi.nlm.nih.gov/entrez/query.fcgi?cmd=Retrieve&db=PubMed&dopt=Citation&list_uids=23750255.

Traynelis, S. F. et al. (2010) ‘Glutamate receptor ion channels: structure, regulation, and function’, Pharmacol Rev, 62(3), pp. 405–496. Available at: http://www.ncbi.nlm.nih.gov/entrez/query.fcgi?cmd=Retrieve&db=PubMed&dopt=Citation&list_uids=20716669.

Valbuena, S. and Lerma, J. (2016) ‘Non-canonical Signaling, the Hidden Life of Ligand-Gated Ion Channels’, Neuron. Elsevier Inc., 92(2), pp. 316–329. doi: 10.1016/j.neuron.2016.10.016.

Vissel, B. et al. (2001) ‘A use-dependent tyrosine dephosphorylation of NMDA receptors is independent of ion flux’, Nat Neurosci, 4(6), pp. 587–596. Available at: http://www.ncbi.nlm.nih.gov/entrez/query.fcgi?cmd=Retrieve&db=PubMed&dopt=Citation&list_uids=11369939.

Wang, Q. et al. (2021) ‘Positive feedback between ROS and cis-axis of PIASx α / p38 α - SUMOylation / MK2 facilitates gastric cancer metastasis’, CDD press, 12(986), pp. 1–12. doi: 10.1038/s41419-021-04302-6.

Wang, Y. et al. (1997) ‘AMPA receptor-mediated regulation of a Gi-protein in cortical neurons’, Natureature, 389(3 October), pp. 502–504.

Waxman, E. A. and Lynch, D. R. (2005) ‘N-Methyl-D-aspartate Receptor Subtype Mediated Bidirectional Control of p38 Mitogen-activated Protein Kinase *’, Journal of Biological Chemistry. Â© 2005 ASBMB. Currently published by Elsevier Inc; originally published by American Society for Biochemistry and Molecular Biology., 280(32), pp. 29322–29333. doi: 10.1074/jbc.M502080200.

Weilinger, N. L. et al. (2016) ‘Metabotropic NMDA receptor signaling couples Src family kinases to pannexin-1 during excitotoxicity’, Nature Neuroscience, 19(3), pp. 432–442. doi: 10.1038/nn.4236.

Wurdemann, T. et al. (2016) ‘Stereotactic injection of cerebrospinal fluid from anti-NMDA receptor encephalitis into rat dentate gyrus impairs NMDA receptor function’, Brain research, 1633, pp. 10–18. doi: 10.1016/j.brainres.2015.12.027.

Zakrzewicz, A. et al. (2017) ‘Canonical and Novel Non-Canonical Cholinergic Agonists Inhibit ATP-Induced Release of Monocytic Interleukin-1 β via Different Combinations of Nicotinic Acetylcholine Receptor Subunits α 7, α 9 and α 10’, Frontiers in Cellular Neuroscience, 11(July), pp. 1–16. doi: 10.3389/fncel.2017.00189.

Zhang, Q. et al. (2012) ‘Suppression of synaptic plasticity by cerebrospinal fluid from anti-NMDA receptor encephalitis patients’, Neurobiology of Disease. Elsevier Inc., 45(1), pp. 610–615. doi: 10.1016/j.nbd.2011.09.019.

