## Supplementary figures and images for "Anti-NMDAR and non-anti-NMDAR antibodies promptly modulate NMDAR through p38 and flux-independent signaling: implications for anti-NMDAR encephalitis"

### Supp fig 1

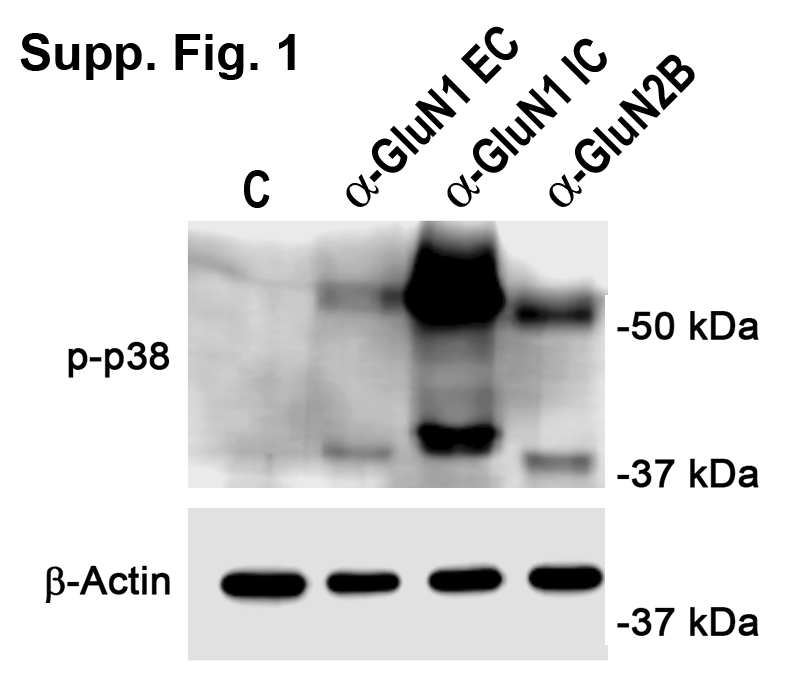

### Supp fig 2

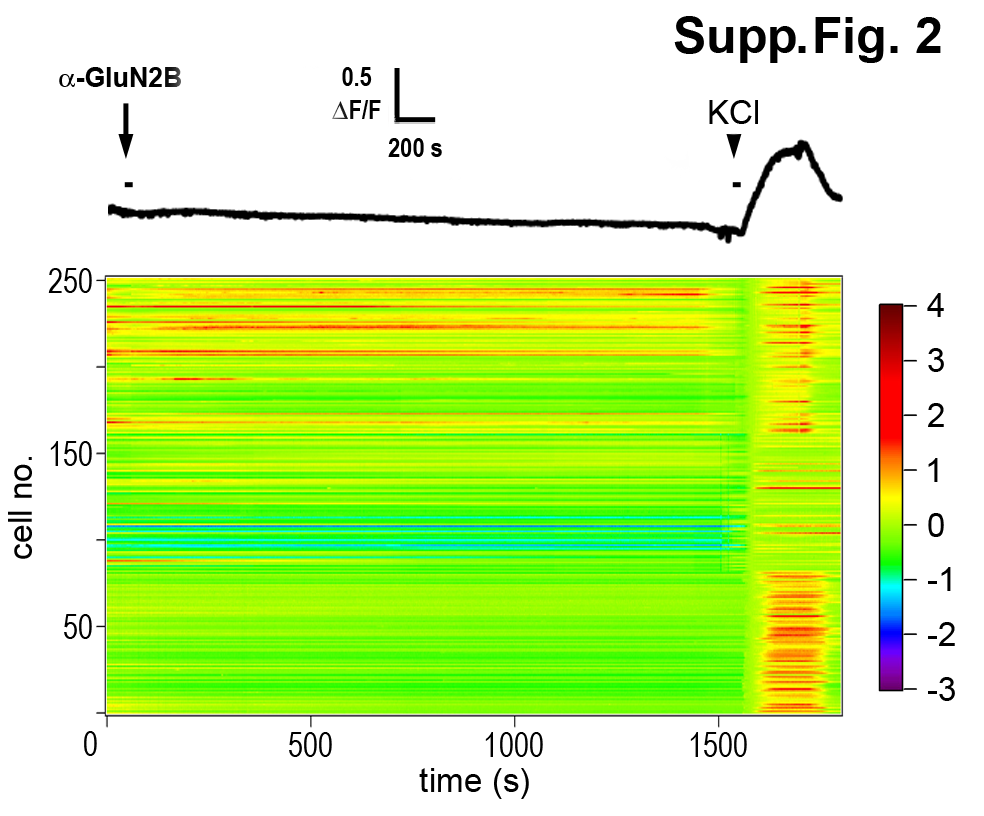

### Supp Table 1

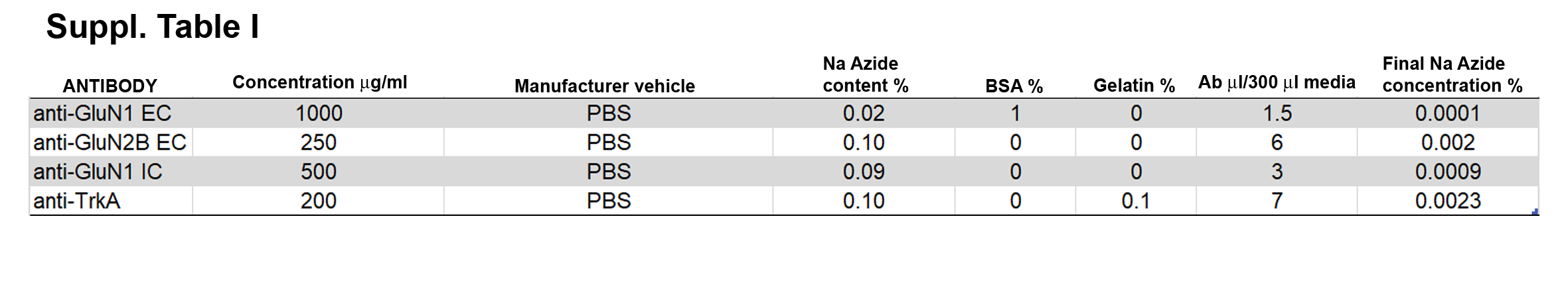
